# Faster carbon accumulation in global forest soils

**DOI:** 10.1101/149393

**Authors:** Weixin Zhang, Yuanqi Chen, Leilei Shi, Xiaoli Wang, Yongwen Liu, Xingquan Rao, Yongbiao Lin, Yuanhu Shao, Xiaobo Li, Shengjie Liu, Shilong Piao, Weixing Zhu, Xiaoming Zou, Shenglei Fu

## Abstract

Comparing soil organic carbon (SOC) stocks across space and time is a fundamental issue in global ecology. However, the conventional approach fails to determine SOC stock in an equivalent volume of mineral-soil, and therefore, SOC stock changes can be under- or overestimates if soils swell or shrink during forest development or degradation. Here, we propose to estimate SOC stock as the product of mineral-soil mass in an equivalent mineral-soil volume and SOC concentration expressed as g C Kg^-1^ mineral-soil. This method enables researchers to compare SOC stocks across space and time. Our results show an unaccounted SOC accumulation of 2.4 - 10.1 g C m^-2^ year^-1^ in the 1m surface mineral-soils in global forests. This unaccounted SOC amounts to an additional C sink of 0.12 – 0.25 Pg C year^-1^, which equals 30 – 62% of the previously estimated annual SOC accumulation in global forests. This finding suggests that forest soils are stronger C sinks than previously recognized.

## INTRODUCTION

Whether a given terrestrial soil functions as a sink or source of atmospheric carbon (C) depends on a precise quantification of the stock and accumulation rate of soil organic carbon (SOC) (Dixon *et al*. 1994; Richter *et al*. 1999; Jobbágy & Jackson 2000; Lal 2004; Stockmann *et al*. 2013). However, there are still large uncertainties in the estimation of SOC accumulation rate which hampers reliable assessments of the response and feedback of terrestrial ecosystems to global changes. The stock of SOC is conventionally calculated by multiplying soil mass with SOC concentration (Adams 1973; Brimhall *et al*. 1991) and summing up to a fixed soil depth, typically 1 m (Pan *et al*. 2011). Then, changes in SOC stocks are estimated across space or over time. However, during soil development or degradation, soil volume for a defined soil mass can either increase (expansion) or decrease (contraction), but seldom stays unchanged. The conventional approach fails to define the total soil mass because it ignores changes in soil volume (ΔV). Consequently, major problems arise when the conventional method for calculating SOC stock (Post & Kwon 2000; Jandl *et al*. 2014; Schuur *et al*. 2015) is used to compare SOC across space or over time (Table 1). Since soil porosity (SP) and soil organic matter (SOM) content influence soil volume as forests develop (Zhou *et al*. 2006; Zou *et al*. 2010), the conventional method will likely underestimate SOC stocks and SOC accumulation rates for soil in developing forests with expanding soil volume. The conventional method can be expressed as:
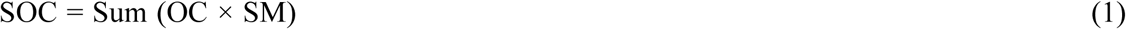

where SOC is SOC stock or density (g C m^-2^), SM is soil mass (g-SM m^-2^), OC is SOC concentration (g C g^-1^-SM), and the Sum function refers to the soil layers added up to a defined soil depth H (typically, H = 1 m).

**Table 1.**
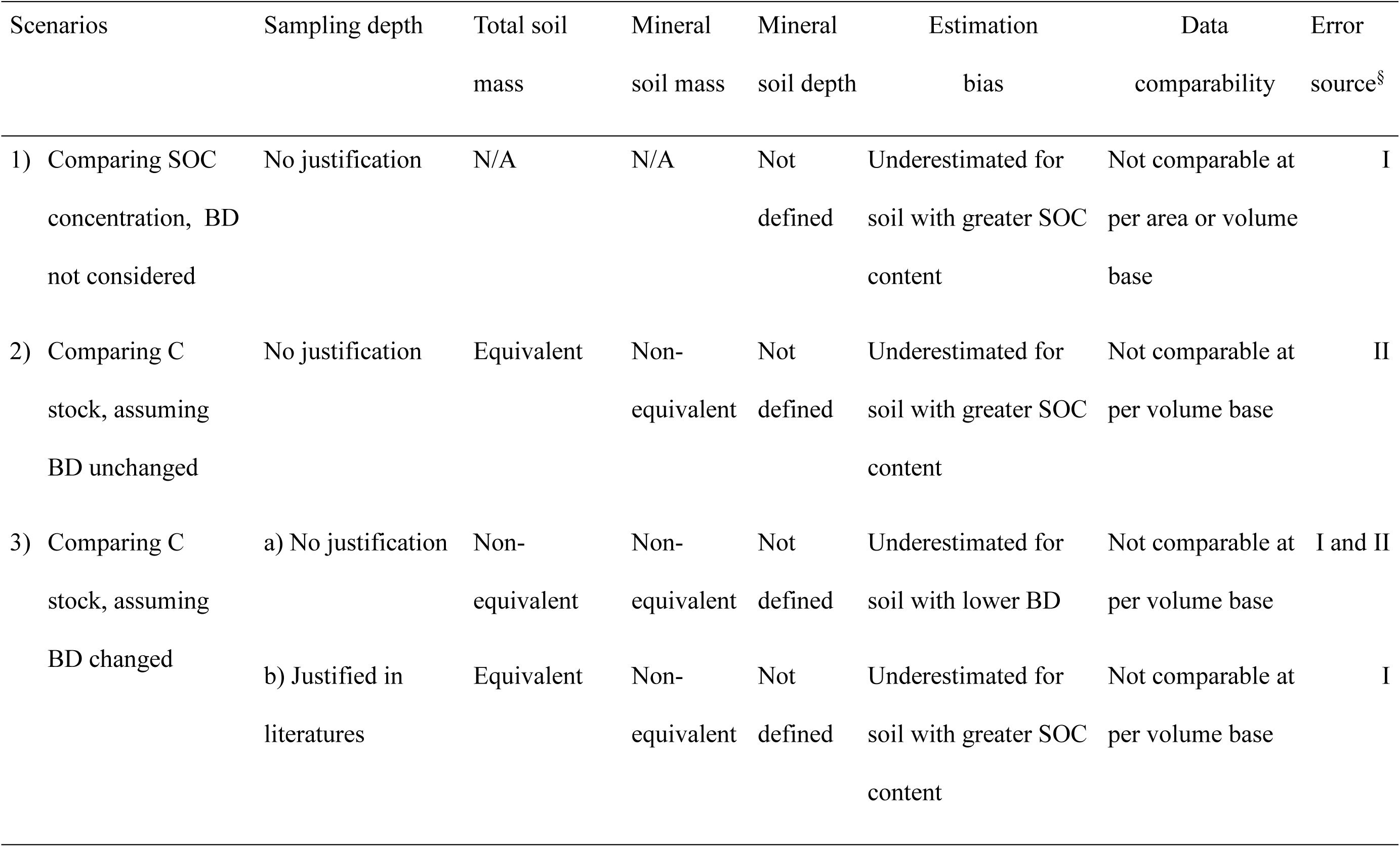
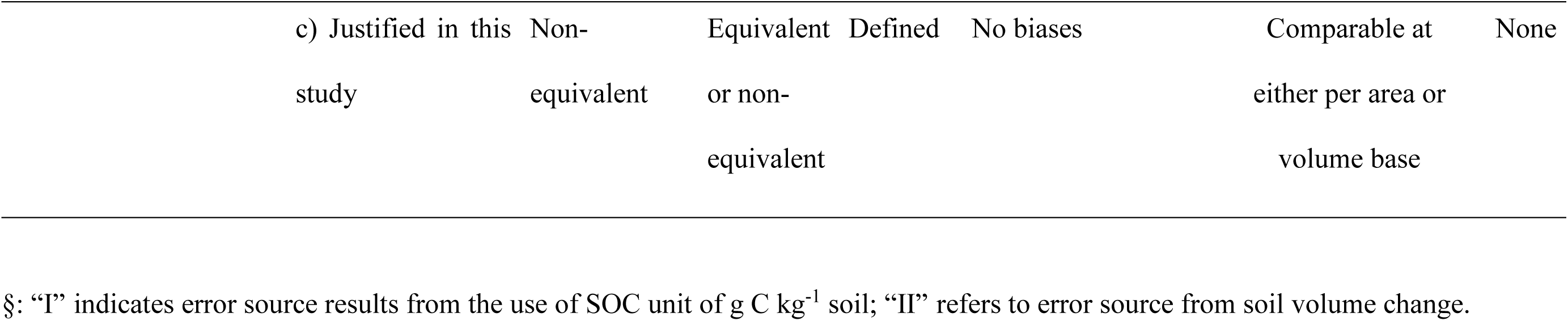
Current methods for soil C stock estimation and their biases.

The conventional method for measuring SOC accumulation rate over a temporal scale between t_1_ and t_2_ only requires calculating SOCt_1_ and SOCt_2_ at the fixed soil depth of H (i.e., Ht_1_ = Ht_2_) and regarding the differences between SOCt_1_ and SOCt_2_ as the SOC accumulation rate during the period. However, SMt_1_ and SMt_2_ may differ due to changes in soil volume resulting from inconsistent SP and/or SOM content. Since an increase in SP and/or SOM, which typically occurs during soil development, will likely reduce the total SM within the fixed soil depth H [i.e., Sum (SMt_1_) > Sum (SMt_2_)], the conventional method underestimates SOCt_2_ and SOC accumulation rate (Fig. 1).

**Fig. 1.**
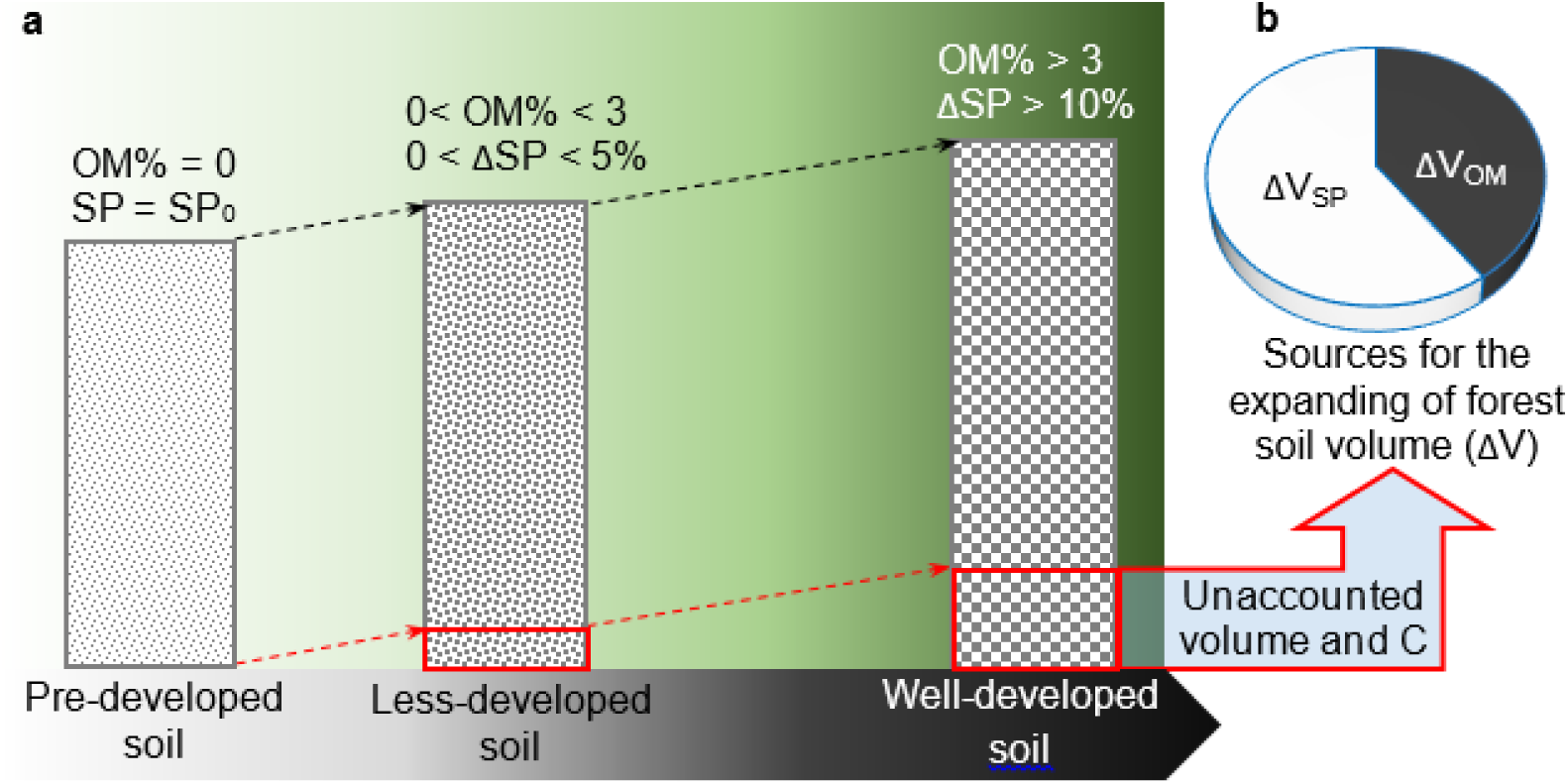
A conceptual framework for the estimation biases of SOC accumulation during forest development. Panel **a** shows how the unaccounted soil volume and C increase with the changes of soil organic matter (SOM) content and soil porosity (SP). Forest and soil development is indicated by the gradation of green and black, respectively. The distance between the black and red arrow lines refers to the soil sampling depth by the conventional approach. Panel **b** shows the sources of soil volume change: changes in SOM and SP. SP_0_: SP in reference soil; ΔSP: change of SP relative to SP0, which consist of the increased SP-occupied volume within mineral soil and SOM; ΔV_OM_: the true volume of SOM (excluding SP volume within SOM).

To overcome this problem of changing soil volume, researchers used an approach of equivalent soil mass (namely the ESM approach) to compare SOC across space and time in several studies (Dalal & Mayer 1986; Ellert & Bettany 1995; Mikhailova *et al*. 2000; Lee *et al*. 2009). In these improved calculations, soil mass (SM) is the same and SMt_1_ equals to SMt_2_, but soil sampling depths do not need to be equal. Nevertheless, this improvement ignores changes in mineral-soil mass (MSM) caused by changing SOM. An increase in SOM will reduce the amount of MSM included in the calculation [i.e. Sum (MSMt_1_) > Sum (MSMt_2_)], resulting in an underestimate of SOCt_2_ and the SOC accumulation rate. Furthermore, even if the influence of SOM on MSM is negligible, the ESM approach generates SOC data at different mineral-soil mass due to inconsistent SP (Table 1). Thus, an alternative approach is to compare SOC in an equivalent mineral-soil mass (EMSM) (Tremblay et al. 2006; Poulton et al. 2003). However, to compare SOC stocks and accumulation rates across space and time, the most reliable and applicable approach is to make comparisons on an equivalent depth basis of mineral-soil so that to avoiding biases induced by inconsistent SP and mineralogical density (Poulton et al. 2003). There are still no studies that successfully bring this thought into operation in SOC accumulation comparisons at different spatial or temporal scales.

Here, we propose a new method to estimate SOC stocks and accumulation rates based on an equivalent mineral-soil volume approach (namely the EMSV approach):
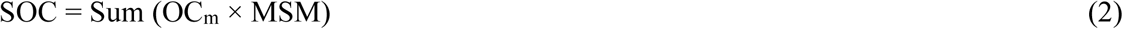

where MSM is the mineral-soil mass in equivalent volume of basal mineral-soil (g-MSM m^-2^); the basal mineral-soil is defined as pre-developed mineral-soil with natural porosity and without organic matter; OCm is the SOC concentration based on MSM (g C g^-1^-MSM), and the Sum function refers to the added MSM up to a defined EMSV for any time [i.e., Sum (MSVt_1_) = (MSVt_2_) = EMSV] and space [i.e., Sum (MSVs_1_) = (MSVs_2_) = EMSV]. We consider all soils in a particular soil type starting from a pre-developed soil (Fig. 1a) and use the pre-developed soil as a reference system to calculate the soil volume change (ΔV) of a given forest soil (Fig. 1b), which will quantify SOC stocks in the defined EMSV (see *Materials and Methods*). Using this method, we then examined the patterns of soil ΔV change and the associated unaccounted C in global forest soils using a compiled global database of forest soil properties (GFSP, Fig. S1 and Supplementary Data). Finally, we re-estimated forest SOC accumulation rates on both local and global scales with this new approach using the GFSP database and literature data from studies of SOC accumulation in forests (Pan *et al*. 2011).

## MATERIALS AND METHODS

### General configuration

In order to compare SOC across space and time in the equivalent mineral-soil volume (EMSV), we needed to first define the MSV (e.g., 1 m depth of basal mineral-soil) and quantify the SOC in the same defined EMSV. We introduced the concept of using pre-developed soils as a standard reference system practically defined as soils without apparent soil developing processes (typically beneath the B horizon). We used this reference system to quantify changes in soil volume (ΔV) resulting from changes in SP (ΔV_SP_) and/or in OM content (ΔV_OM_, Fig. 1b). We quantified the unaccounted MSM and the associated SOC and, then, recalculated the SOC stocks (Modified C_density_) and accumulation rates (Modified *K*C_density_). The ΔV for a given soil profile with a fixed sampling depth at a given time was calculated by comparing the volumes derived from soil porosity (SP) and organic matter (OM) with those in the reference pre-developed soil profiles. Accordingly, based on equation 2, the unaccounted SOC (ΔC_density_) was calculated as a product of mineral soil mass (MSM) and SOC concentration (OC_m_) in the expanded soil horizon. In this way, SOC accumulation rates over a given time interval can be calculated, and compared across space and time. The major part of a given MSV has been accounted for in the conventionally sampled soil layers using equation 1, but a proportion of MSV may be unaccounted for due to soil volume expansion. Therefore, the total SOC stock in a given MSV is the sum of the accounted and unaccounted SOC. In other words, we considered that the conventional sampling depth is not enough for any given soil sample to keep the defined EMSV. The new method we proposed successfully includes the unaccounted mineral soil mass so that the comparison of SOC across space and time is applicable. The main equations are as below:
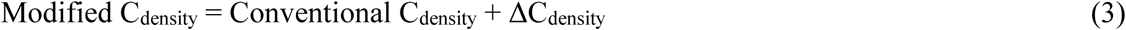

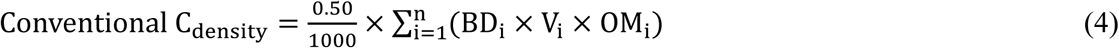

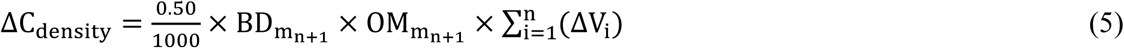

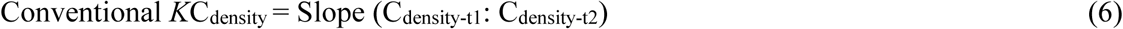

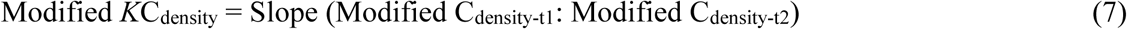

where Modified C_density_ and Conventional C_density_ refers to SOC stock estimated by the conventional method and our modified method, respectively (g C m^-2^ soil); ΔC_density_ refers to the unaccounted SOC stock (g C m^-2^ soil) for a given sampled volume of soil if comparing SOC in EMSV; BD is g soil cm^-3^ soil; V is the sampled soil volume for a given soil horizon, cm^3^ m^-2^; OM is the organic matter concentration, g OM kg^-1^ soil; “i” refers to the number of soil horizon for a given soil profile; 0.50 is the conversion factor from OM to C (Pribyl 2010). BD_m_ is the BD of mineral soil (g mineral soil cm^-3^ soil); OM_m_ is the organic matter associated with each unit of mineral soil (g OM kg^-1^ mineral soil); ΔV_i_ refers to the ΔV of the “i”th horizon in the profile (cm^3^ m^-2^); “n” refers to the last (deepest) soil horizon for a given soil profile; “n + 1” refers to the adjacent deeper soil horizon with volume of ΔV (the total soil volume change for a given profile); and BDm_n+1_ and OMm_n+1_ refer to BD_m_ and OM_m_ in the expanded soil horizon, respectively; Conventional *K*C_density_ refers to the SOC accumulation rate in a given non-equivalent soil mass (NESM) during a given time interval (g C m^-2^ year^-1^); Modified *K*C_density_ is the SOC accumulation rate in a given EMSV during a given time interval (g C m^-2^ year^-1^); t1 and t2 refers to the start and end times of a given duration.

### The database of global forest soil properties (GFSP)

In order to estimate the global patterns of OM, SP, SP_0_, BD and the annual relative change in BD (RCBD, g cm^-3^ year^-1^), we established a database for global forest soil properties (GFSP), which consists of 961 plots, and 4184 rows of data (Appendix S1; Fig. S1; Supplementary Data).

### Estimation of soil volume change

The volume increases of OM (ΔV_OM_) and SP (ΔV_SP_) are two major sources of soil volume change (ΔV). They can be estimated by comparing OM and SP in the reference soil (OM = 0; SP = SP_0_) with those in the studied soils. The main equations are below:
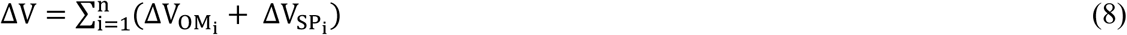

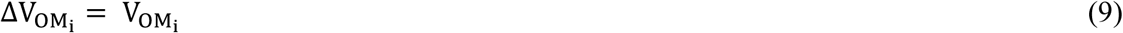

If the soil profile contains several horizons (n > 1), V_OM_ and ΔV_SP_ in the “i”th horizon (i <= n - 1) can be calculated as:
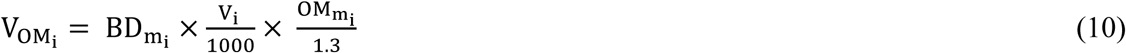

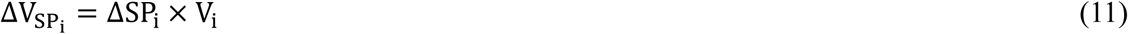

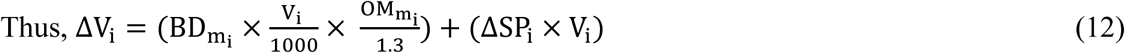

For the last soil horizons (i = n and n > 1) or if the soil profile contains only one horizon (n = 1), V_OM_ and ΔV_SP_ in the “n” horizon can be calculated as:
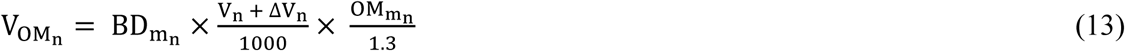

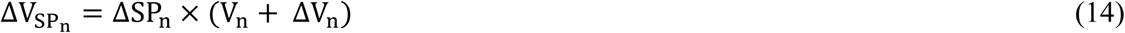

Thus,

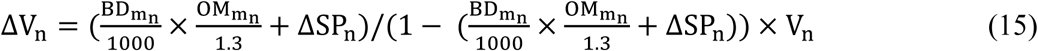

Here,
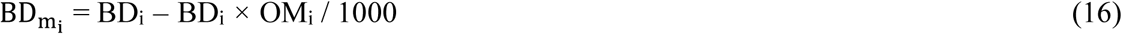

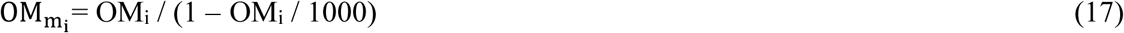

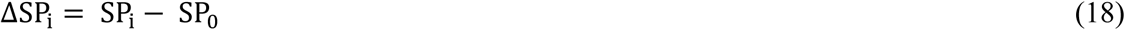

Where ΔV refers to the total soil volume change for a sampled soil profile; “i” refers to the number of soil horizon in the soil profile; BD_m_ is the BD of mineral soil (g mineral soil cm^-3^ soil); V_i_ is the sampled soil volume of the “i”th horizon, cm^3^ m^-2^; OM_m_ refers to the organic matter content (g OM kg^-1^ mineral soil); ΔV_i_ is the soil volume change in the “i”th horizon; 1.3 is the true density of OM (g cm^-3^) (Adams 1973); SP_0_ refers to the soil porosity in the pre-developed soil, which is estimated from the averages of the minimum values of SP (excluding any values > 60%) of the deep soil layers (> 40 cm) in each plot for a given biome using the database of GFSP. Interestingly, the estimated SP_0_ does not differ across biomes (*F*_2,271_ = 1.02, *P* = 0.363; Fig. S2), suggesting that a common SP_0_ (43.9%) can be used in estimating ΔV_SP_ and ΔV_OM_ at the global scale.

If SP is not given, it can be calculated from soil BD and OM. We made an improvement to the conventional equations for both BD and SP (Adams 1973; Post & Kwon 2000). We partitioned soil volume into three components: (a) true volume of OM (V_OM_, excluding porosity within OM), (b) true volume of mineral soils (V_M_, excluding porosity within mineral particles), and (c) the total volume of soil porosity within both OM and mineral particles (V_SP_). Thereby, we introduce new equations for BD and SP as below:
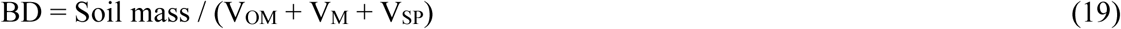

thus for 100 g of soil sample, the equation can be rephrased as:
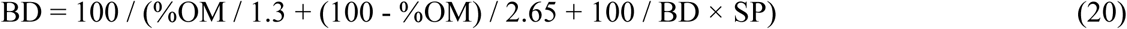

and further rephrased as: 
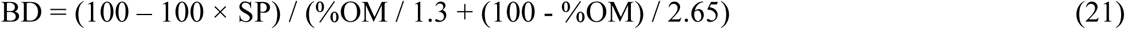

and accordingly, SP can be calculated as:
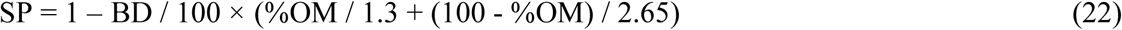

where %OM is per cent by weight of OM; 2.65 is the true density of mineral soils (Post & Kwon 2000). Note that when %OM is zero, our equation for SP is equal to the conventional SP equation (i.e., SP = 1 – BD / 2.65) indicating that the conventional equation for SP is not suitable for soil with high content of OM.

### Calculation of unaccounted C stock in the 10 cm standardized forest soil horizons

We classified the database of GFSP into three biomes, three OM levels, and three soil layers., and normalized the depth of all soil horizons to 10 cm (Appendix S2). Given that the standardized 10 cm soil horizon is considered as an independent uniform unit (Fig. S3), we assumed that BD_m_ and OM_m_ in the expanded part of soil were equal to those in the 10 cm soil horizons. Thus, to calculate the Modified C_density_ and ΔC_density_, the equations 3 - 5 can be simplified as: 
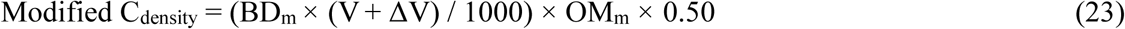

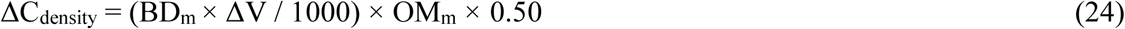

### Calculation of unaccounted forest SOC stock in the whole soil profiles

In order to represent forest soil profiles across biomes, six forest sites with various climate and soil characteristics were selected from the GFSP database (Appendix S3). The unaccounted SOC stocks in the whole profiles with varied depths were calculated using our modified method. For each plot (one site may have several plots), a linear and an exponential model was established to describe the BD and OM as a function of soil depth (h, cm), respectively (Table S1). Thereby, BDm_n+1_ and OMm_n+1_ were calculated with those equations, where “h” equals the original depth of a given soil profile plus its expanded depth (Δh). Afterwards, the unaccounted C for each soil profile, plot, and site were estimated by equation 5.

### New estimation of forest SOC accumulation rate at a long-term study site

In order to quantify the method-derived uncertainty in SOC accumulation rate, we re-analyzed the soil C dataset from an old-growth monsoon evergreen forest at Dinghushan Mountain where soil C was found to accumulate with stand age (Zhou et *al*. 2006). We used the two equations (SOC = 0.035x – 67.97, R^2^ = 0.90, *P* < 0.0001 and BD = -0.0032x + 7.42, R^2^ = 0.90, *P* = 0.01; here “x” refers to years) in Fig. 1 of Zhou *et al*. (2006) to calculate the SOC (%) and soil BD in the surface layer (0 – 20 cm) from 1979 to 2003. Based on equation 4, the conventional C_density_ for a given year (C_density-year_) was calculated, and based on equation 6, the C_density_ change rate from the conventional method (Conventional *K*C_density_, g C m^-2^ year^-1^) was calculated as:
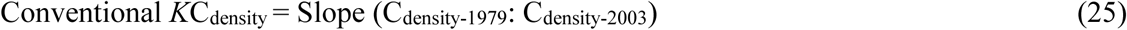

In addition, based on equations 8 and 13 - 18, we calculated BD_m_, OM_m_, ΔSP, and ΔV for the surface soil profile (0 – 20 cm). Furthermore, we calculated the decreasing rates of C_density_ (*S*C_density_) with depth (i.e., from 0 – 20 cm to 20 – 40 cm) with literature data (Table S2) (Fang *et al*. 2003; Zhang 2011). Then, the unaccounted C stock (ΔC_density_, g C m^-2^) for a given year from 1979 to 2003 could be estimated as:
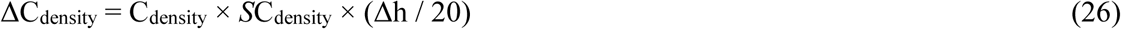

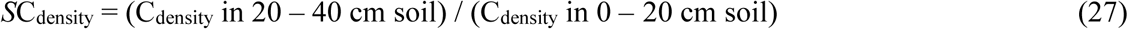

Finally, the modified SOC stock for a given year was calculated with equation 3, and the modified SOC accumulation rate (Modified *K*C_density_, g C m^-2^ year^-1^) was estimated as:
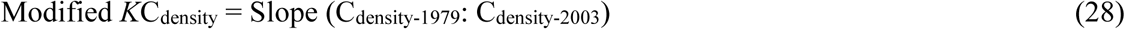

### New estimation of global forest SOC accumulation rate

We re-analyzed the dataset of global forest SOC dynamics from 1990 to 2007 in Pan *et al*. (2011). This re-calculation did not include sites in Japan and South Korea since no data were available. In addition, we noticed that the total forest C density (including C in both living/dead plant biomass and soil) declined from 1990 to 2007 in temperate Europe and New Zealand. These trends may imply that the forest qualities in these regions are declining and, thus, soil volume change may be limited. Therefore, these two regions were also excluded in this study to reach a more conservative estimate of global forest SOC accumulation rates. Given that the dataset in Pan *et al*. (2011) only showed forest soil C density (Mg C ha^-1^) and total forest area (Mha) in 1990, 2000, and 2007, we needed to first give an initial value of soil BD (i.e., determine BD value in 1990) based on the GFSP-derived mean forest BD (Table S3). Then, the annual relative changes in bulk density (RCBD) of forest soils across biomes were calculated (Appendix 4) using the GFSP database and data from Zhou *et al*. (2006) (Table S4). The equation is:
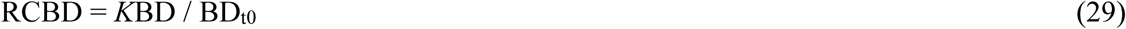

where, *K*BD is the slope of soil BD and BD_t0_ refers to soil BD at time zero (i.e., the beginning of a specific forward development of forests). Finally, the values of global forest soil BD in 2000 and 2007 were calculated based on the given values of BD in 1990 and the RCBD. Thereby, soil OM, OM_m_, BD_m_, and SP in 1990, 2000, and 2007 were calculated using the dataset of forest SOC density (g C m^-2^) in Pan *et al*. (2011) and the given/calculated BD. Then, we re-calculated the global forest soil C density (C_density_, g C m^-2^) during 1990 to 2007 with our modified method and compared with them to those derived from the conventional method. Forest SOC accumulation rates (*K*C_density_, g C m^-2^ year^-1^) and the change rates of total forest SOC stock (*K*C_stock_, Pg C year^-1^) for a given region are calculated as below: 
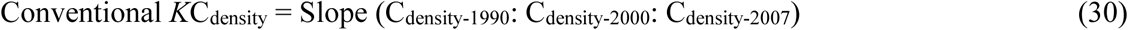

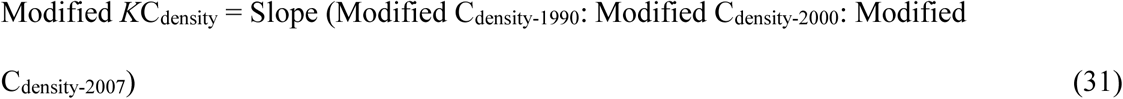

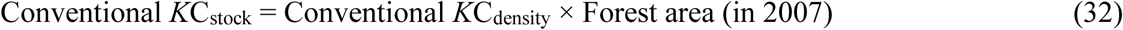

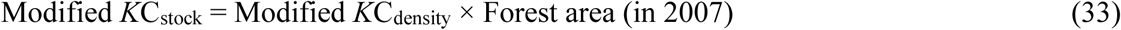

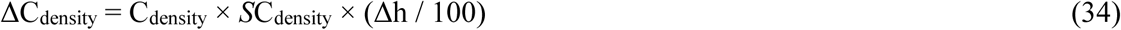

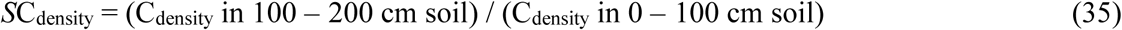

Note that the maximum and minimum values of the decrease rates of C_density_ (*S*C_density_) with depth (i.e., from 0 – 100 cm to 100 – 200 cm) in different biomes were calculated with literature data (Table S5) (Jobbágy & Jackson 2000), thus, the ranges of unaccounted forest SOC were also estimated. Given that C density in the upper portion of a soil horizon is normally greater than that in the lower portion, our approaches (Equation 26 and 34) will underestimate ΔC_density_; thus, the forest SOC accumulation rates are likely still underestimated. Here, the unaccounted SOC includes a fraction of OC that is not included in the mineral soil mass of the initial 1 m soil, but this should not significantly contribute to more unaccounted C because soil volume changes and their contribution to the estimation bias of SOC stock mainly occur in the surface soils. We focused on the 1 m mineral soil mass equivalent depth so that SOC stocks could be compared across space and time.

Additionally, in order to exclude forest lands with limited expansion of soil volume, we estimated the proportions (*f*) of forest plots in which SP was lower than the reference soil porosity (SP_0_) (Table S6) using the database of GFSP. Thus, the more conservative estimation of global forest SOC accumulation rate (*K’*C_stock_, Pg C year^-1^) was calculated as: 
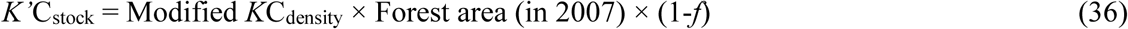

### Statistical methods

One-way ANOVA was performed to compare the SP_0_ and the unaccounted SOC accumulation rate among different biomes, and to examine the effect of the sampling depth of soil profiles on the amount of unaccounted SOC stocks from six representative forest sites across biomes. Either the post hoc LSD test (for homogeneous variances) or Tamhane’s T_2_ test (for non-homogeneous variances) was performed for multiple comparisons. General linear model was used to test the main effects of biome, soil layer and OM level on the amounts and proportion of unaccounted SOC stock in the GFSP-derived standardized 10 cm forest soil horizons. All statistics were performed with SPSS 19.0.

## RESULTS

### Theoretical Patterns of ΔV and Unaccounted SOC in Global Forests

To illustrate the differences in the conventional method with fixed soil depth and our modified method with fixed mineral soil mass, we calculated and compared soil volumes and SOC stocks (C density) in the standardized 10 cm soils and 10 cm mineral soils using the GFSP-derived dataset. The SOC stocks calculated in the standardized 10 cm soils with traditional method ranged from 913 to 7682 g C m^-2^ in boreal forests, 549 to 5807 g C m^-2^ in temperate forests, and 687 to 6106 g C m^-2^ in tropical forests (Table S7). The postulated pre-developed 10 cm mineral soils expanded 0.28 - 4.33 cm, 0.53 - 4.72 cm, and 0.75 - 6.18 cm in boreal, temperate, and tropical forests, respectively (Table S7). In boreal, temperate, and tropical forests, the increase in volume of OM contributed to 1.4 - 11.8%, 0.8 - 8.9%, and 1.1 - 9.4% of the soil volume expansions, respectively, and the increase volume in SP contributed to 1.2 - 17.9%, 3.7 - 25.1%, and 5.5 - 28.8% of the soil volume expansions (Table S7).

The corresponding unaccounted SOC stocks were 27 - 3084 g C m^-2^, 39 - 2507 g C m^-2^, and 63 - 3776 g C ^m-2^ (Fig. 2a-c) and accounted for 2.8 - 43.3%, 5.3 - 47.1%, and 7.5 - 61.8% of the SOC stocks calculated by the conventional method for the respective boreal, temperate, and tropical forests (Fig. 2d-f). The SOM level and biome type had significant impacts on both the amount (*F* = 635.5, *P* < 0.001 and *F* = 4.74, *P* = 0.009, respectively) and proportion (*F* = 236.2, *P* = 0.000 and *F* = 11.9, *P* = 0.000, respectively) of unaccounted forest SOC. Soil layer only significantly affected the amount of unaccounted SOC (*F* = 3.28, *P* = 0.038).

**Fig. 2.**
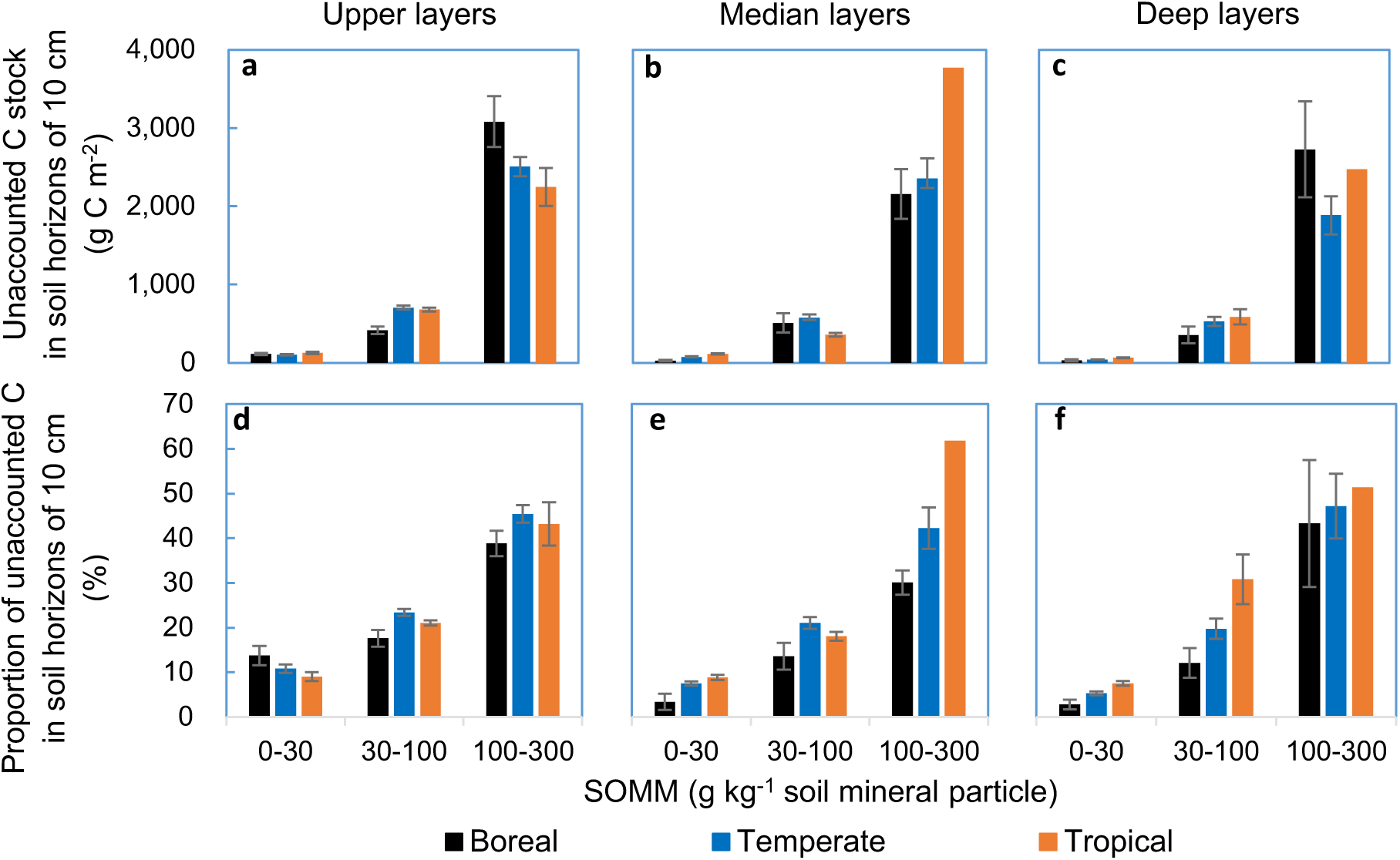
Global patterns of the amount (**a-c**) and proportion (**d-f**) of unaccounted SOC stock in standardized 10 cm mineral soil. Mean values in the Upper (0 − 20 cm), Median (20 – 40 cm), and Deep soil layers (> 40 cm) are shown ± 1 s.e.m., as a function of SOM content and biome types.

### Unaccounted Forest SOC Stocks in the Whole Soil Profile

To characterize the changes in ΔV and unaccounted C in the whole soil profile, we calculated the stock of unaccounted forest SOC in soil profiles ranging in depth from 5 – 60 cm. We selected six representative forest sites across biomes from the GFSP-derived dataset. On average, the unaccounted forest SOC stocks in soil profiles with varied depths were 1035 - 5083 g C m^-2^, 899 - 1043 g C m^-2^, and 630 - 1040 g C m^-2^ in boreal, temperate, and tropical forest sites, respectively (Fig. 3). Unexpectedly, for most soil profiles, the amount of unaccounted SOC does not decrease significantly with an increase in sampling depth (Fig. 3).

**Fig. 3.**
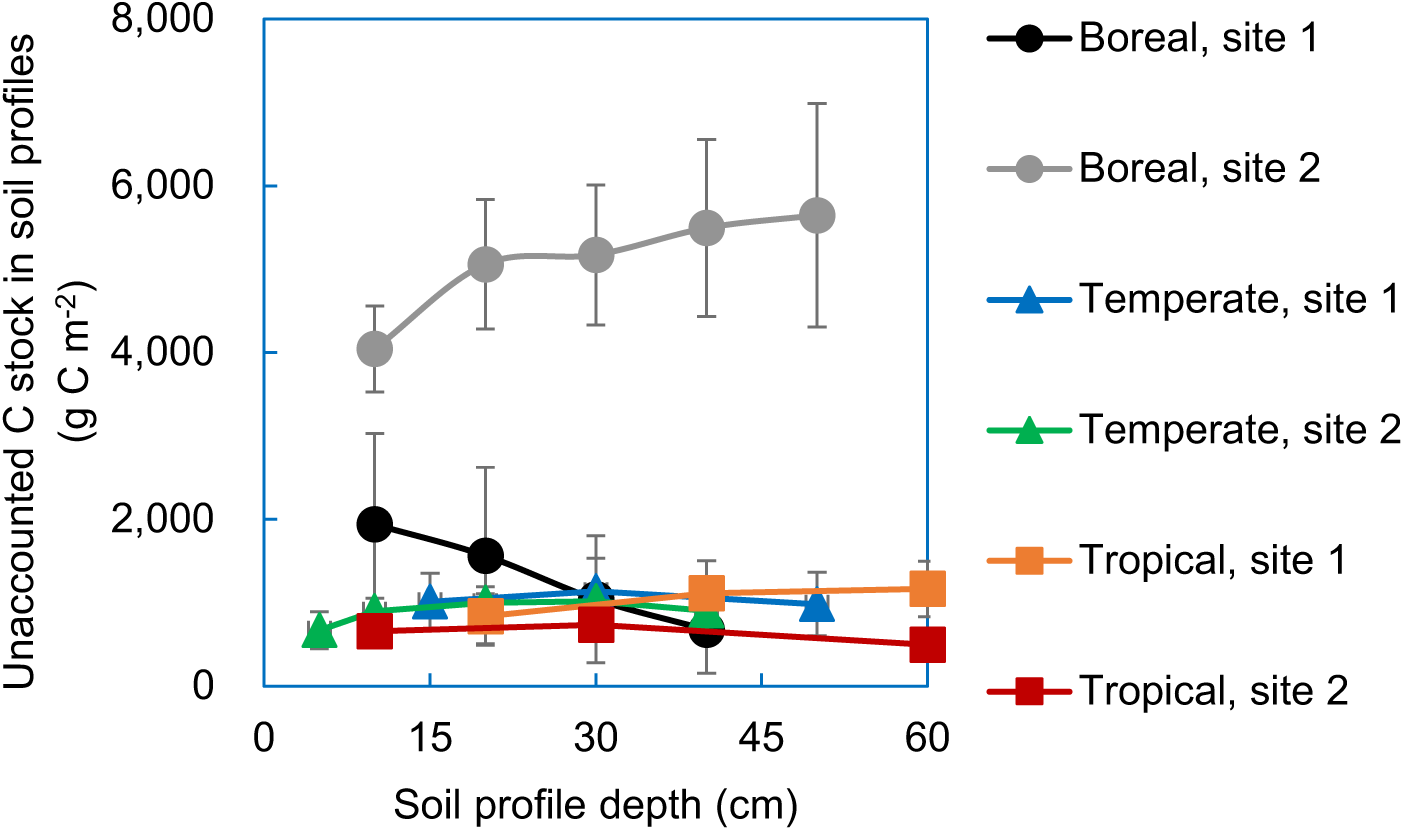
Unaccounted SOC stock in soil profiles along soil depth for representative forests across biomes. Mean values are shown ± 1 s.e.m.; the effects of sampling depth on the amounts of unaccounted C in soil profiles are tested: Boreal site 1: Amuer (*F*_3,8_ = 0.393, *P* = 0.762), Boreal site 2: Tianlaochi (*F*_4,25_ = 0.440, *P* = 0.779); Temperate site 1: Mao county (*F*_2,6_ = 0.046, *P* = 0.955), Temperate site 2: Sanming (*F*_4,5_ = 1.032, *P* = 0.473); Tropical site 1: Pingxiang (*F*_2,3_ = 0.237, *P* = 0.802) and Tropical site 2: Jianfengling Mountain (*F*_2,359_ = 30.95, *P* = 0.000).

### Unaccounted Forest SOC Accumulation Rate in a Local Mature Forest

To show the estimation biases in forest SOC change over time, we re-calculated SOC accumulation rate (0 - 20 cm depth) in a well-studied tropical old-growth forest where SOM concentration and bulk density (BD) have been monitored over 25 years (Zhou *et al*. 2006). The re-calculated SOC accumulation rate is 13.5% higher than that derived from the conventional method, and only 0.2% lower than the previously assumed upper bound (estimated based on the assumption of a constant BD during forest development) (Fig. 4).

**Fig. 4.**
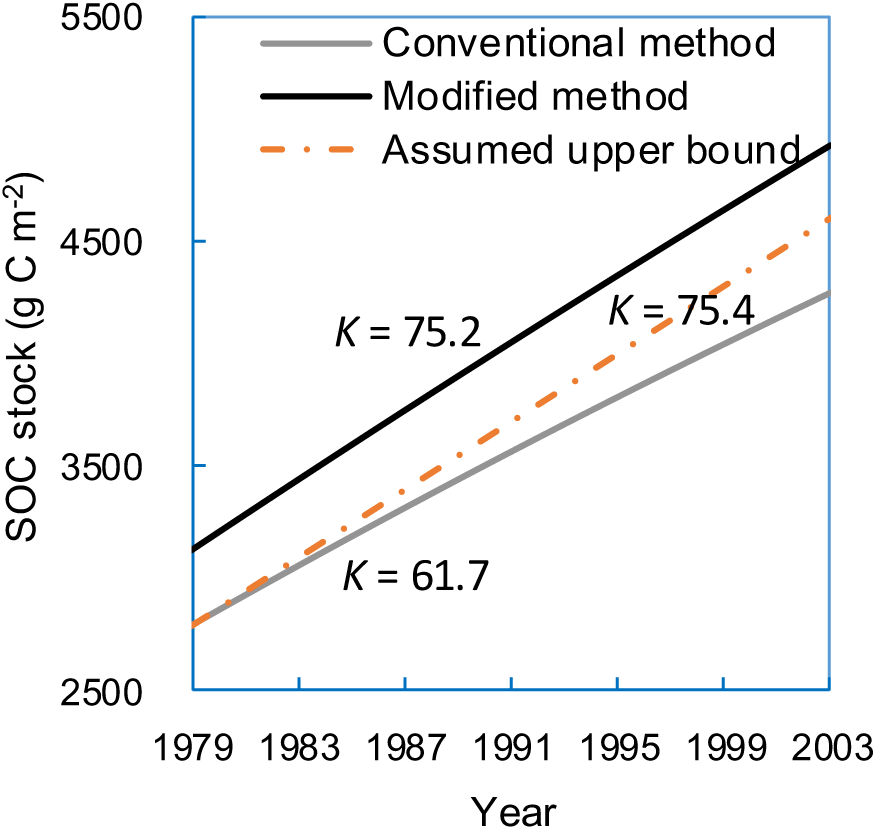
Re-estimated annual SOC accumulation rate in an old-growth forest. The SOC accumulation rates, indicating as line slopes (*K*), were re-calculated by both the conventional method and our modified method using data from the old-growth tropical forests in Dinghushan Mountain (Zhou *et al.* 2006). The line of assumed upper bound refers to C accumulation rate that estimated based on the assumption of a constant BD during forest development.

### Unaccounted SOC Accumulation Rate in Global Forests

Finally, to show the unaccounted forest SOC accumulation rates globally, we re-analyzed a published global dataset of forest SOC to a depth of 1 m (Pan *et al*. 2011). We found that the unaccounted forest SOC sinks in the 14 major forest regions (Pan *et al*. 2011) ranged from 0.001 to 0.089 Pg C year^-1^ (Fig. 5a). From boreal forests to tropical forests, the unaccounted annual SOC accumulations range from 2.4 ± 0.3 to 10.1 ± 0.8 g C m^-2^ year^-1^ (Fig. 5b). The accumulation rates in the boreal forests and the tropical intact forests were greater than those in the temperate forests (low-bound: *P* < 0.001 and *P* = 0.016, respectively; upper-bound: *P* = 0.001 and *P* = 0.005, respectively). Overall, in addition to the previously estimated global forest SOC sink of 0.4 Pg C year^-1^ using the conventional method, we found an additional forest SOC sink of 0.15 - 0.32 Pg C year^-1^ during 1990 - 2007 (Fig. 5c) (Pan *et al*. 2011). The boreal forests and the tropical intact forests contributed to 40 - 48% and 32 - 37% of the global unaccounted forest SOC sink, respectively. Notably, our new calculation indicates that the tropical intact forest soils in the Americas and South Asia are actually important C sinks (3.8 - 9.2 and 2.4 - 7.1 g C m^-2^ year^-1^, respectively) instead of C sources as previously reported (-0.06 and -1.0 g C m^-2^ year^-1^, respectively) (Pan *et al*. 2011).

**Fig. 5.**
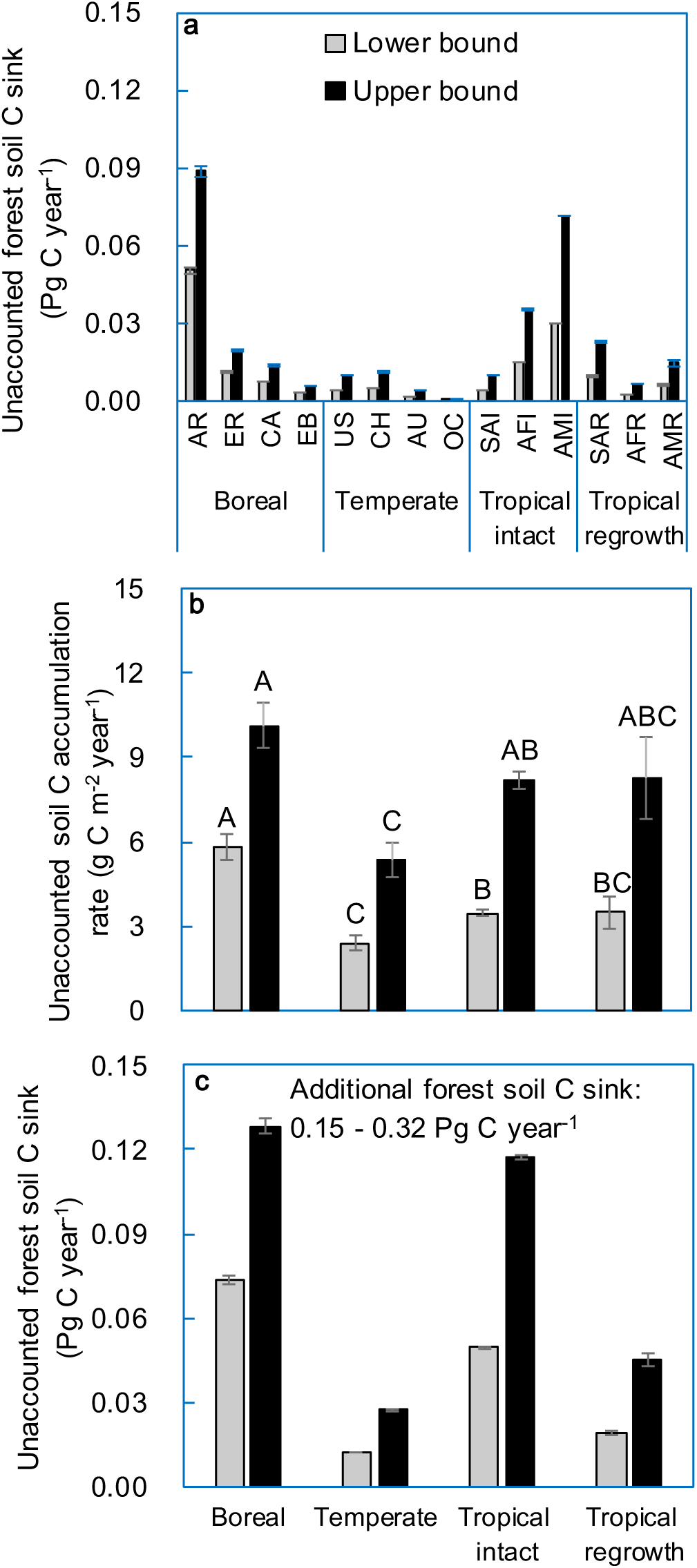
Unaccounted forest SOC accumulation at regional (**a**) and global (**b, c**) scales. In all panels, mean values are shown ±1 s.e.m. Panels **a** and **b** show the re-estimated forest SOC accumulation rate in 14 major regions (AR: Asian Russia, ER: European Russia, CA: Canada, EB: European boreal; US: United States, CH: China, AU: Australia, OC: Other countries in temperate; SAI: South Asia intact, AFI: Africa intact, AMI: Americas intact, South Asia regrowth, AFI: Africa regrowth, AMI: Americas regrowth) and three biomes (Pan *et al*. 2011); bars with different lowercase and uppercase letters indicate significant differences of lower bound and upper bound SOC accumulations, respectively (*P* < 0.05). Panel **c** shows the unaccounted annual global forest SOC sink relative to recently reported values (Pan *et al*. 2011), assuming constant soil volume change over time.

## DISCUSSION

Comparing SOC stocks across space and time is a fundamental issue in global ecology. However, the conventional method of calculating SOC stocks fails to account for the soil volume changes during soil development. This intrinsic flaw creates great uncertainty in the estimation of forest SOC accumulation rate.

Our new approach addresses this problem by calculating SOC stock as the product of mineral-soil mass in equivalent basal mineral-soil volume (EMSV) and SOC concentration expressed as g C Kg^-1^ mineral-soil (OC_m_). First, we created the reference forest soil profiles for tracking and comparing C dynamics using a defined EMSV baseline. Second, we used the EMSV-based approach to calculate forest SOC stocks in standardized 10 cm soil layers from a global forest soil dataset. This work illustrated how changes in OM and SP were connected with changes in soil volume and consequently changes in SOC stocks across different biomes. The results suggested that SOM level is the most important factor that positively affects the unaccounted forest SOC stocks due to its influence on both the OM-occupied and SP-occupied volumes.

Then, we used the EMSV-based approach to show the unaccounted forest SOC stocks in the whole soil profile. The unaccounted forest SOC stocks in soil profiles in boreal forests seemed to be greater than those in temperate and tropical forest sites. Such pattern was probably due to the greater SOC level in boreal forests. It is worthy to note that the unaccounted C for whole soil profiles did not change markedly with sampling depth, especially when depth was > 30 cm. Theoretically, the unaccounted C for the whole soil profile is determined by two parts: (a) the magnitude of ΔV in the sampled soil column, and (b) the SOC concentration beneath the sampled soil column. Since soil volume expansion occurred to the greatest extent in the surface soils (e.g., 0 - 30 cm depth), most of the soil ΔV can be included when surface soils are sampled. Furthermore, SOC concentration beneath surface soils declined rapidly with soil depth and was usually in consistently low levels (Table S1; Jobbágy & Jackson 2000). As a result, major changes in ΔV and SOC concentration are likely to be accounted for as long as the surface soils are included. Therefore, we suggest that it may be sufficient to measure surface soils (e.g., 0 - 30 cm depth) to quantify the total amount of unaccounted SOC. This reduces a tremendous amount of field effort to quantify and compare local and global SOC accumulation rates because most available data are derived from the surface soil layers.

Finally, using this new approach, we re-estimated forest SOC accumulation over time both locally, using a forest in southern China, and globally. After more than 20 years of field monitoring, researchers found that the old-growth forest in Dinghushan Mountain in subtropical China continuously accumulated SOC (Zhou *et al*. 2006). However, our new calculations suggest that this mature forest is even a stronger C sink than previously estimated. On the global scale, our new calculations suggest that the conventional method profoundly underestimates the capacity of C sequestrations in forest soils and sometimes makes an incorrect judgment on whether an ecosystem is a C sink or source.

Nevertheless, there are still some uncertainties in our modified method. On the one hand, the reference soil porosity (SP_0_) used in this study (43.9%) may be greater than the SP in a specific forest site. This may result in an underestimation of soil volume expansion and, therefore, an underestimation of forest SOC accumulation. On the other hand, since a greater proportion of forest lands may be in the process of degradation with soil volumes shrinking over time, the present global C sink in forest soils may be overestimated. To explore these uncertainties, we estimated the proportion of forest lands that do not have significant soil volume expansion using the GFSP database. On average, 21.4% of forest lands show no soil volume expansion relative to the reference soil profiles (Table S6). Consequently, a more conservative estimate of the unaccounted global forest SOC accumulation would be 0.12 - 0.25 Pg C year^-1^. Therefore, the total global C accumulation in forest soils from 1990 - 2007 is 0.52 - 0.65 Pg C year^-1^, which is 30 - 62% greater than that calculated by the conventional method (Pan *et al*. 2011).

Deforestation and land-use changes for agriculture often decrease levels of SOC and SP (Post & Kwon 2000; Murty *et al*. 2002; Li *et al*. 2015). Using the conventional method to measure the effects of these anthropogenic disturbances might result in an overestimation of SOC stock and an underestimation of land use change-induced SOC loss. Our approach using a more comprehensive calculation has the potential to improve the understanding of the impacts of forest development and land use change on the global C budget. To quantify forest SOC accumulation rate, we recommend to first measure SP in several soil samples from deeper layers (e.g., > 100 cm) and use the average value as an approximation of SP0 for a specific site. Secondly, sample surface soil layers (e.g., 0 - 30 cm) between time intervals and calculate the SOC stock and volume change (ΔV) at each time. Finally, sample the next deepest soil layer based on the calculated depth change (Δh, normally < 10 cm for a 30 cm soil profile) and quantify the unaccounted C in the corresponding ΔV soil.

## ACKNOWLEDGEMENTS

This work was supported by the CAS/SAFEA International Partnership Program for Creative Research Teams, the South China Botanical Garden Program for Foster Featured Institute for Young Scholars (Y561011001), the Natural Science Foundation of China (31570516, 41171219), the National Basic Research Program of China (2011CB403205), the NSFC-Guangdong Joint Fund (U1131001), and the Youth Innovation Promotion Association of CAS (2014318). We thank Quan Chen for help to draft the global plot map and Drs. Dan Binkley, David Coleman, Jingyun Fang and Yude Pan for the critical comments on the manuscript.

## SUPPORTING INFORMATION

Additional supporting information is available. The database for global forest soil properties (GFSP) is shown in a separate Excel file.

## Supporting Information

## Appendix S1. The database of global forest soil properties (GFSP)

In order to estimate the global patterns of OM, SP, SP_0_, BD and the annual relative change in BD (RCBD, g cm^-3^ year^-1^), we established a database for global forest soil properties (GFSP). The major data sources included were WISE3, SPADE, and others (Literature search and the National Ecosystem Research Network of China, CERN). We searched in the Web of Knowledge using the key words “bulk density”, “soil porosity”, “bulk density” AND “organic matter”, “soil porosity” AND “organic matter”. Articles including information about either OM, BD, or SP, both OM and BD or SP were collected, and other information such as forest type, age, climate, geographical position and disturbance was recorded. Primary natural forests (occasionally natural shrubs), secondary natural forests, and plantations of more than five years old were included in the database, which consists of 961 plots, and 4184 rows of data (Fig. S1; Supplementary Data). The map of plots was produced in ArcGIS 10.2 with a free basemap from http://www.esri.com/data/find-data.

## Appendix S2. Data preparation for the calculation of unaccounted C stock in the 10 cm standardized forest soil horizons

We classified the database of GFSP into three biomes (boreal, temperate and tropical forests), three OM levels (0 - 30, 30 - 100, and 100 -300 g OM kg^-1^ mineral-soil), and three soil layers (upper layer of around 0 - 20 cm, median layer of around 20 - 40 cm and deep layer of > 40 cm). Generally, we grouped forests with an annual mean temperature of < 0°C, 0 - 20°C and > 20°C into boreal forests, temperate forests and tropical forests, respectively. In order to focus on C dynamics of mineral soil, all surface horizons with an OM level of > 300 g OM kg^-1^ mineral-soil were excluded. Finally, to facilitate comparison among different soil horizons, the depth of all soil horizons was normalized to 10 cm.

## Appendix S3. Forest sites selection for the calculation of unaccounted forest SOC stock in the whole soil profiles

In order to represent forest soil profiles across biomes, six forest sites with various climate and soil characteristics were selected from the GFSP database. The unaccounted SOC stocks in the whole profiles with varied depths were calculated using our modified method. The sits in boreal forests included two sites with contrasting characteristics: 1. Amuer, Daxinganling mountain range, a northern site with a low elevation of 500 - 800 m; 2. Tianlaochi, Heihe River, a relatively southern site with a high elevation of 3100 - 3400 m. The temperate forests sites included one cold temperate forest site (Mao county, Sichuan with an annual mean temperature of 9.3°C, and an elevation of 1785 - 2131 m) and one warm temperate forest site (Sanming, Fujian with an annual mean temperature of 18.8°C). Tropical forests sites included one in Pingxiang, Guangxi with an annual mean temperature of 20.5 - 21.7°C and another in Jianfengling Mountain, Hainan with an annual mean temperature of 25°C. The six sites were all in China because we did not find many available datasets of BD, SP, and OM for the whole soil profile, including several horizons and replications at each plot, from other regions.

## Appendix S4. The calculation of annual relative changes in bulk density (RCBD) in global forest soils

Given that the dataset in Pan *et al*. (2011) only showed forest soil C density (Mg C ha^-1^) and total forest area (Mha) in 1990, 2000, and 2007, we needed to first give an initial value of soil BD (i.e., determine a value of BD in 1990). Our simulations of soil OM content (calculating OM content by giving a range of values of BD) indicated that OM contents in the 1 m deep of soil profiles were within our lowest OM category (i.e., 0 - 30 g C kg^-1^ mineral soil). Hence, the GFSP-derived mean forest BD for soils with low content of OM_m_ (0 - 30 g kg^-1^ mineral soil) was used (Table S3). Since soil BD varies with soil layer for all biomes, we assumed that the given BD values may be a source of uncertainties for the estimations of forest SOC stocks and change rates.

Then, the annual relative changes in bulk density (RCBD) of forest soils across biomes were calculated based on the GFSP database and data from Zhou *et al*. (2006) (Equation 29). Few studies provided RCBD or data that could be used to calculate RCBD (Table S4). The values of RCBD in boreal forests, which were derived from only two studies at one site, were much greater and more variable (-0.0255 ± 0.008 g cm^-3^ year^-1^) than those from the temperate and tropical soils. However, the values of RCBD were very close between the temperate and tropical forests (-0.0036 ± 0.0008 and -0.0030 g cm^-3^ year^-1^, respectively). We then assumed the same values of RCBD in boreal forests as those in temperate forests to make a conservative estimate of soil volume increment rate. Then, the values of soil BD in 2000 and 2007 were calculated based on the given values of BD in 1990 and the RCBD.

**Fig. S1.**
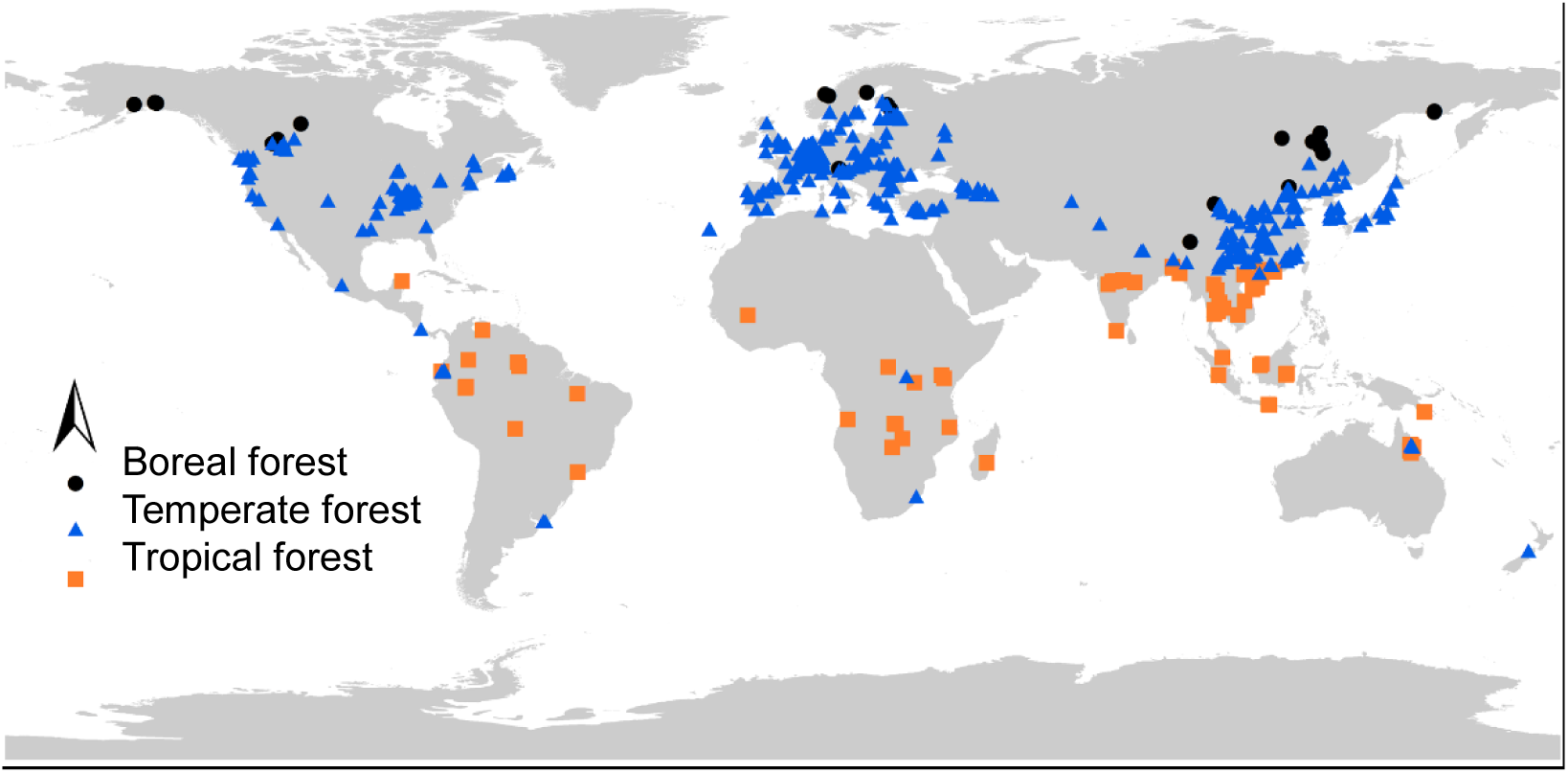
The distribution map of plots for establishing the database of global forest soil properties (GFSP). Primary natural forests (occasionally natural shrubs), secondary natural forests, and plantations of more than five years old are included in the database, which consists of 961 plots (Supplementary Data). The map of plots is produced in ArcGIS 10.2 with a free basemap from http://www.esri.com/data/find-data.

**Fig. S2.**
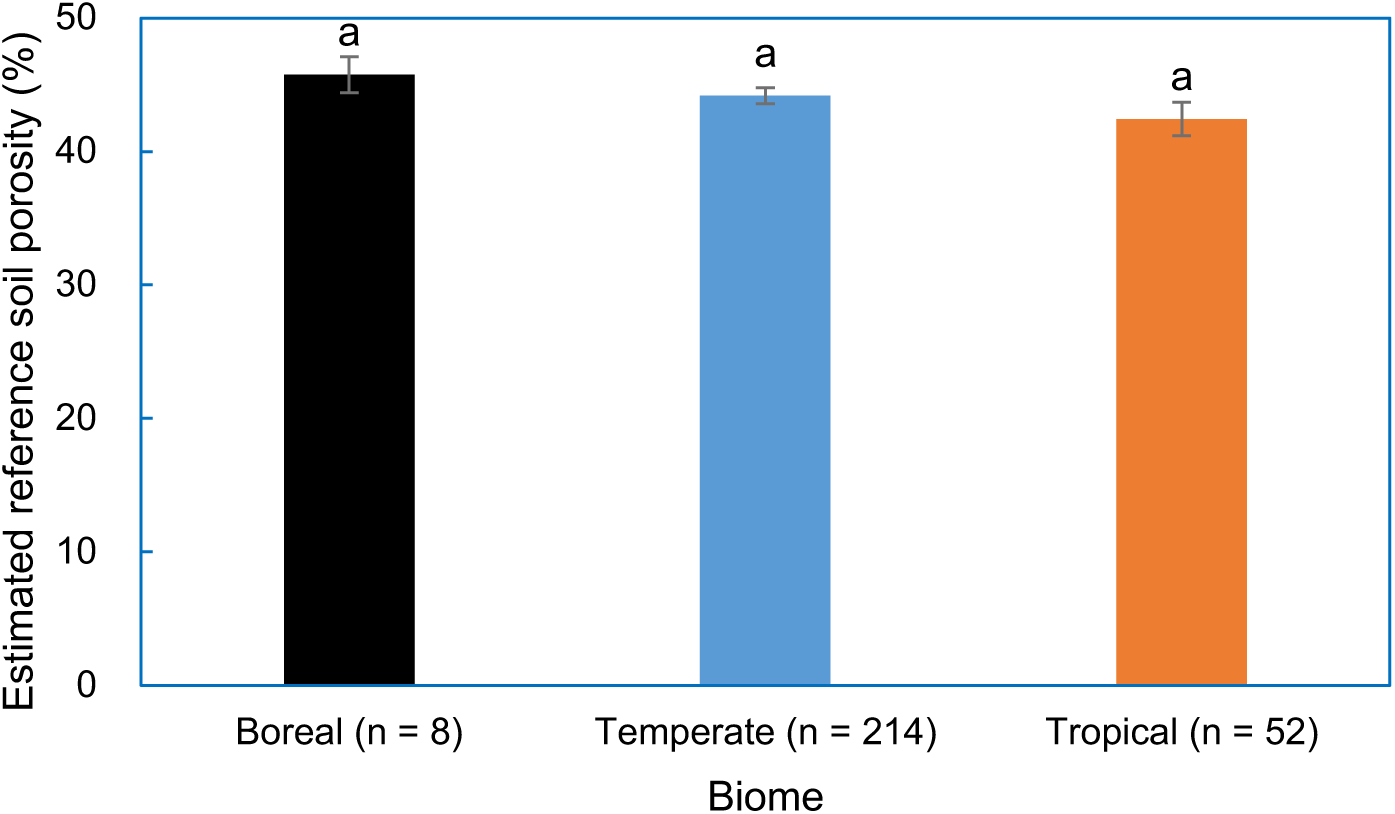
Global pattern of estimated soil porosity in pre-developed forest ecosystems (SP_0_). Mean values are shown ± 1 s.e.m.; bars with same letters indicate non-significant differences of reference soil porosity (*P* > 0.05).

**Fig. S3.**
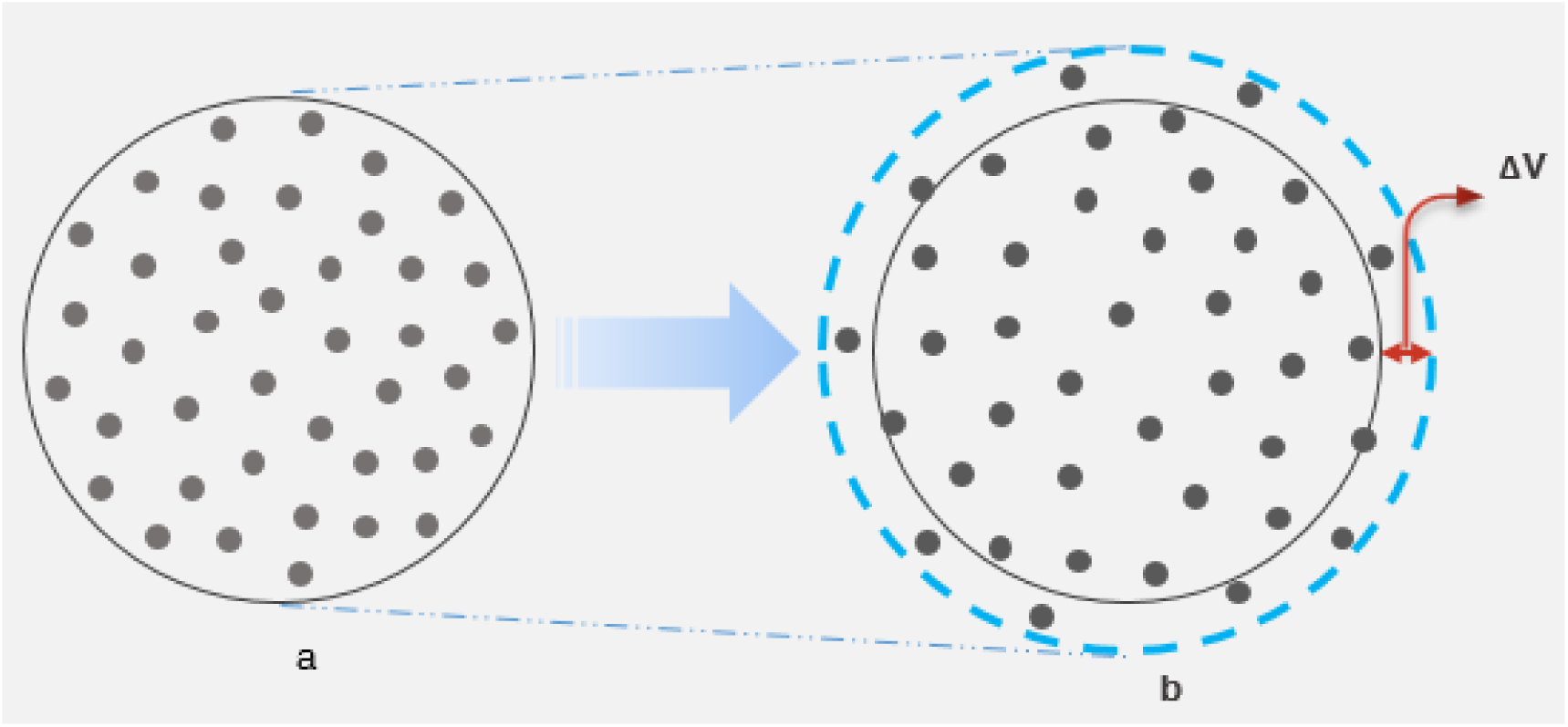
A diagram for soil volume change in a standardized horizon. Panel **a** refers to a pre-developed forest soil horizon with lower soil porosity and negligible amount of organic C; panel **b** refers to a relatively well-developed forest soil horizon with greater soil porosity and higher organic C concentration. The circle dots represent soil mineral particles; darker color means greater C concentration. the same numbers of dots indicate same mass of soil mineral particles between **a** and **b**; the circle dots being located between the solid and dotted lines in panel **b** indicate the unaccounted mineral soil and associated C due to soil volume change (ΔV).

**Table S1.**
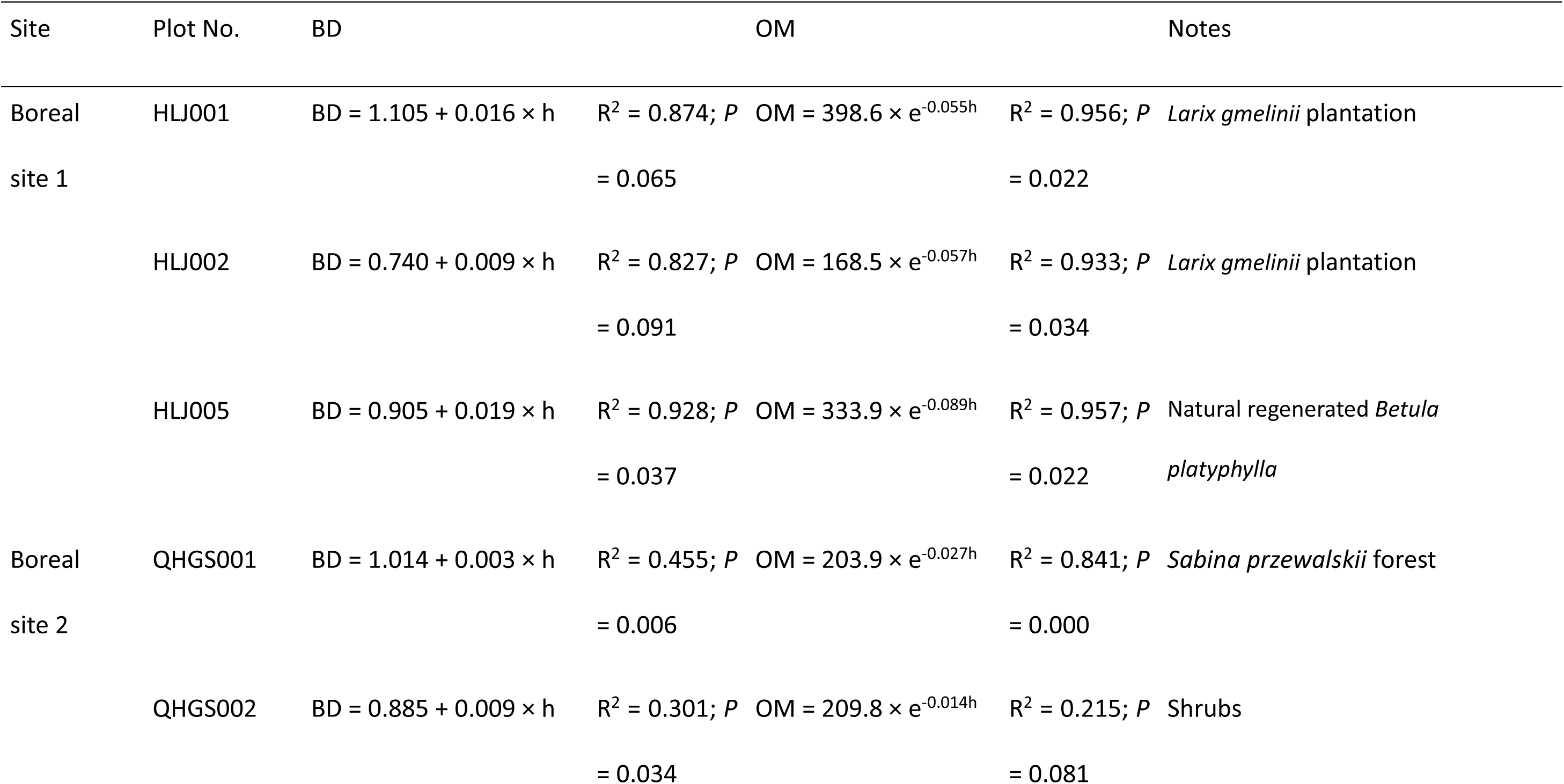
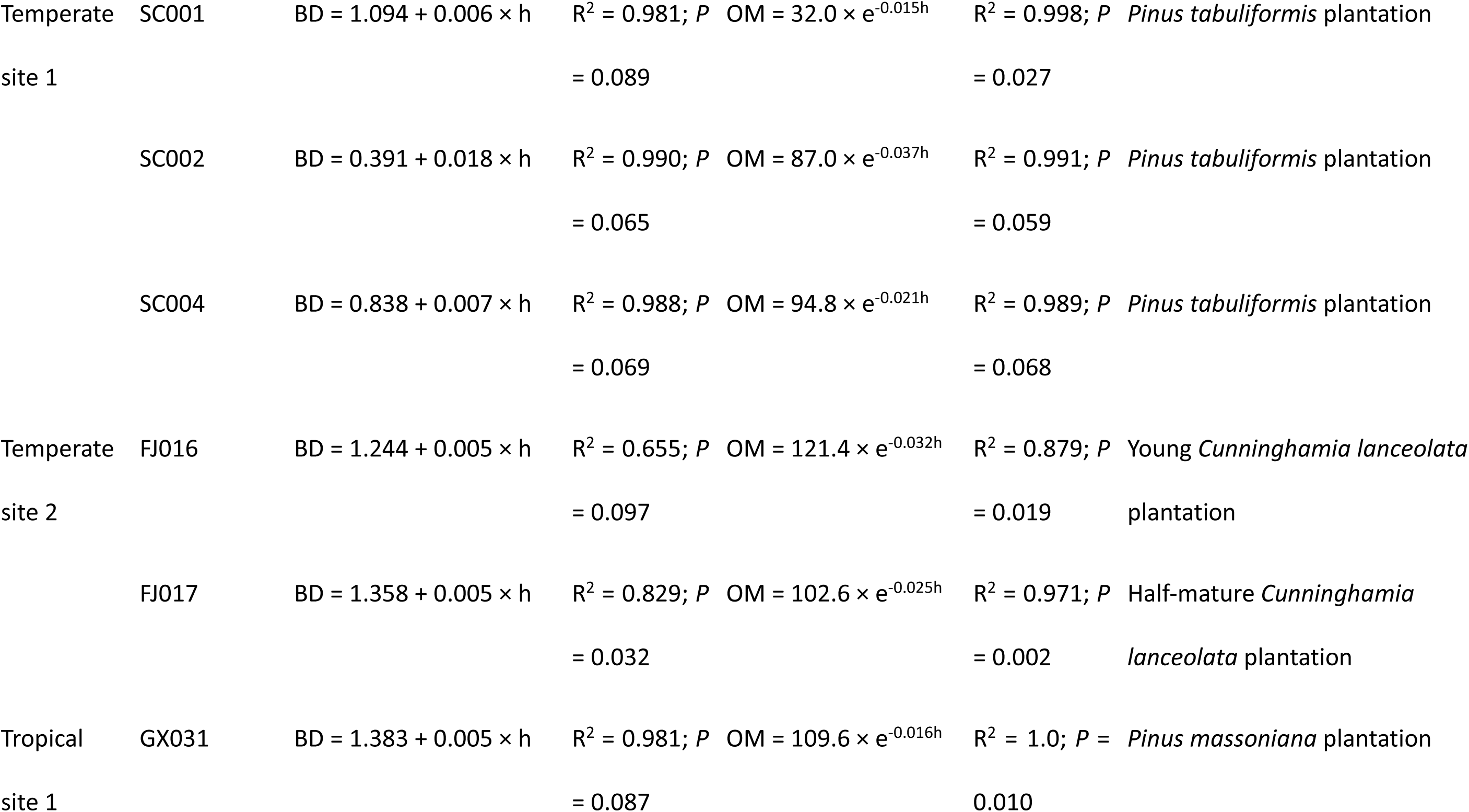
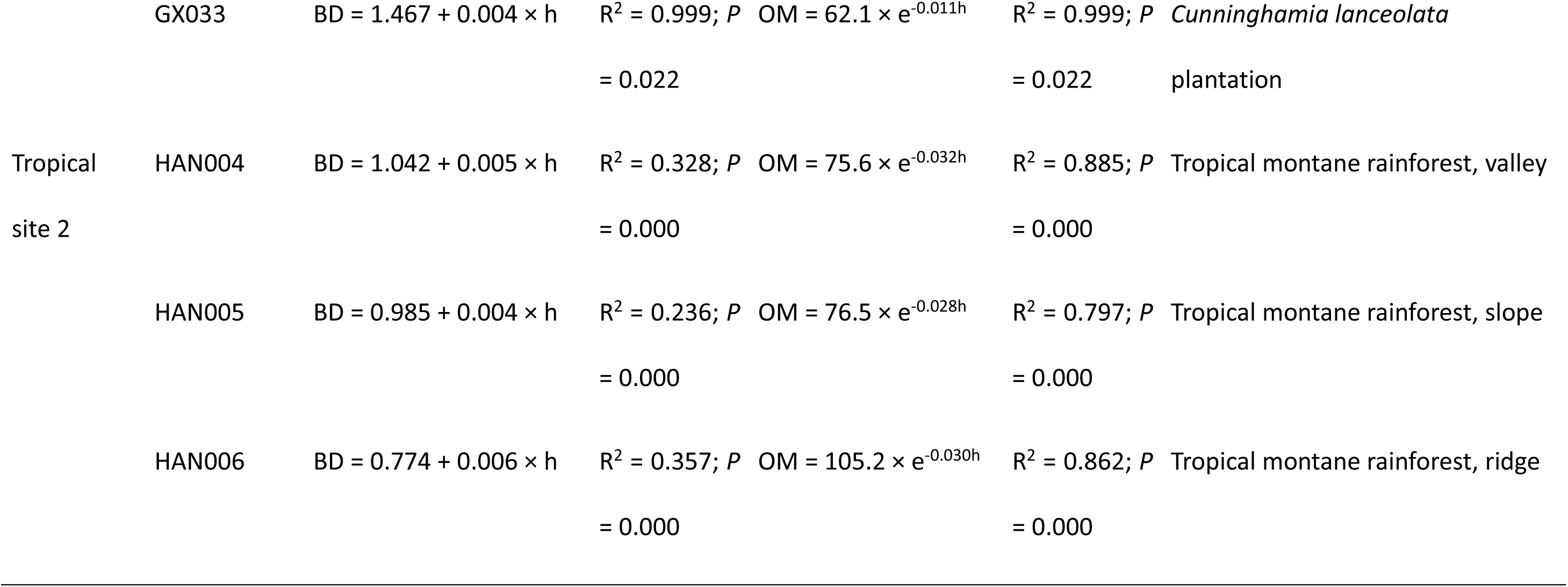
Models describing the variation of BD and OM with soil depth (h, cm) at six representative sites across biomes.

**Table S2.** The decrease rates of C_density_ (*S*C_density_) with depth in the old-growth monsoon evergreen forest at Dinghushan Mountain.

| Soil horizon<br>(cm) | SOC<br>(g C kg<sup>-1</sup> soil) | BD<br>(g cm<sup>-3</sup>) | $C_{\text{density}}$<br>(g C m<sup>-2</sup>) | $SC_{\text{density}}^{\dagger}$ | Reference |
| --- | --- | --- | --- | --- | --- |
| 0 - 10 | 31.4 | 0.936 | 2939 | 0.581 | Zhang 2011 |
| 10 - 20 | 11.9 | 1.276 | 1516 |  |  |
| 20 - 40 | 10.2 | 1.269 | 2591 |  |  |
| 0 - 10 | 32.3 | 0.844 | 2726 | 0.585 | Fang *et al.* 2003 |
| 10 - 20 | 20 | 0.964 | 1928 |  |  |
| 20 - 40 | 12.4 | 1.098 | 2723 |  |  |
| Average |  |  |  | 0.583 |  |

$\dagger SC_{\text{density}} = (C_{\text{density}} \text{ in } 20 - 40 \text{ cm soil}) / (C_{\text{density}} \text{ in } 0 - 20 \text{ cm soil})$

**Table S3.** Mean estimated forest soil BD for soils with low content of OM_m_ (0 - 30 g kg^-1^ mineral soil) across biomes based on the database of GFSP.

| Biome | Soil layer | BD | n |
| --- | --- | --- | --- |
| Boreal | Upper layer | 1.04 ± 0.10 | 23 |
|  | Median layer | 1.39 ± 0.08 | 9 |
|  | Deep layer | 1.46 ± 0.04 | 6 |
| Temperate | Upper layer | 1.30 ± 0.02 | 133 |
|  | Median layer | 1.37 ± 0.01 | 208 |
|  | Deep layer | 1.47 ± 0.01 | 333 |
| Tropical | Upper layer | 1.39 ± 0.03 | 46 |
|  | Median layer | 1.28 ± 0.02 | 107 |
|  | Deep layer | 1.34 ± 0.01 | 246 |

**Table S4.**
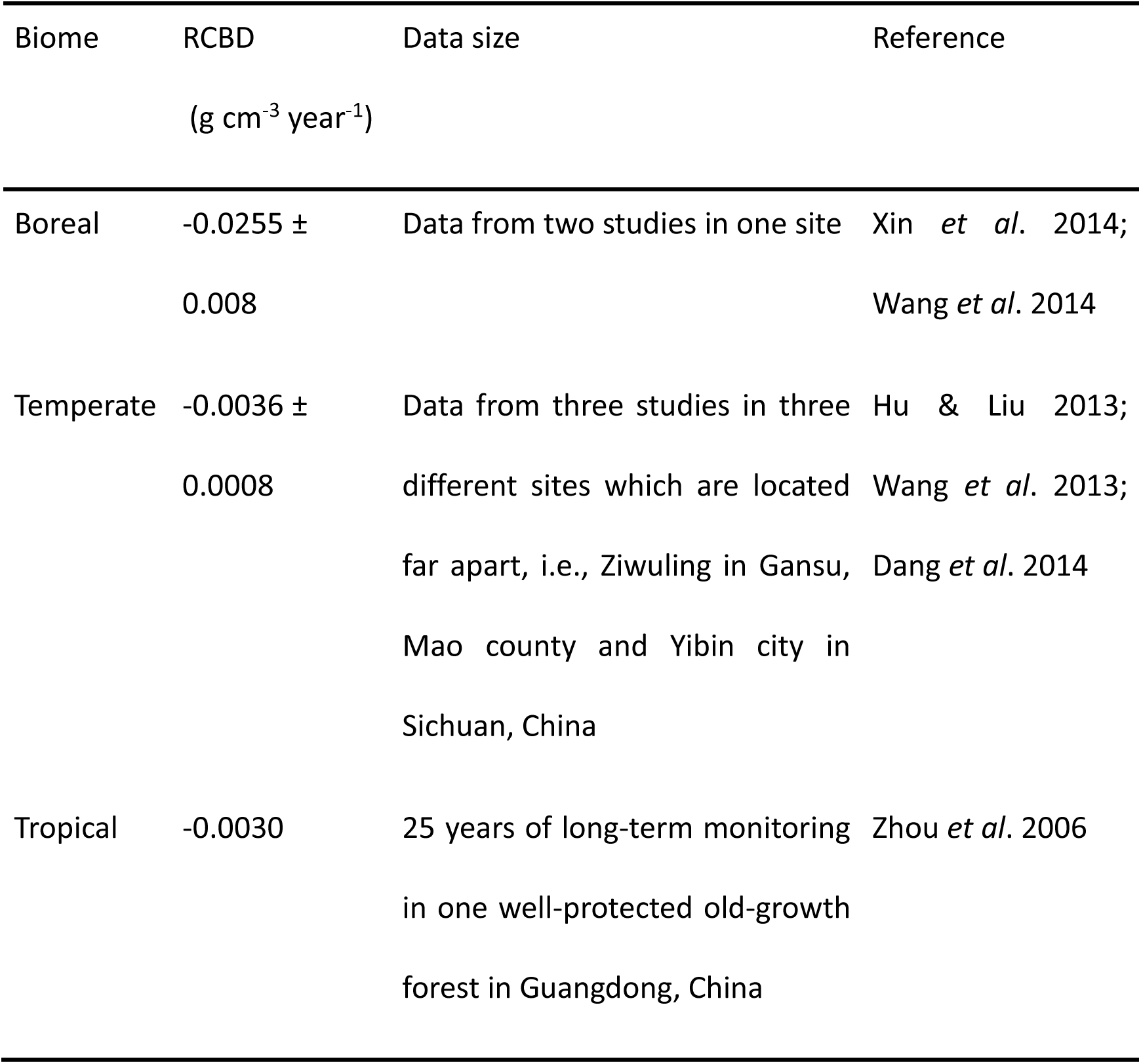
Annual relative changes of bulk density (RCBD) of forest soils across biomes.

| Biome | RCBD<br>(g cm<sup>-3</sup> year<sup>-1</sup>) | Data size | Reference |
| --- | --- | --- | --- |
| Boreal | -0.0255 ± 0.008 | Data from two studies in one site | Xin *et al.* 2014;<br>Wang *et al.* 2014 |
| Temperate | -0.0036 ± 0.0008 | Data from three studies in three different sites which are located far apart, i.e., Ziwuling in Gansu, Mao county and Yibin city in Sichuan, China | Hu & Liu 2013;<br>Wang *et al.* 2013;<br>Dang *et al.* 2014 |
| Tropical | -0.0030 | 25 years of long-term monitoring in one well-protected old-growth forest in Guangdong, China | Zhou *et al.* 2006 |

**Table S5.** The maximum and minimum decrease rates of C_density_ (*S*C_density_) with soil depth in different biomes†.

| Biome | SOC content (kg m<sup>-2</sup>) by depth (cm) |  |  |  |  |  | Maximum | Minimum |
| --- | --- | --- | --- | --- | --- | --- | --- | --- |
|  |  |  |  |  |  |  | SCdensity | SCdensity |
|  | 0 – 100 cm |  |  | 100 – 200 cm |  |  |  |  |
|  | Mean | 95%CI,<br>upper bond | 95%CI,<br>lower bond | Mean | 95%CI,<br>upper<br>bond | 95%CI,<br>lower bond |  |  |
| Boreal forest | 9.3 | 9.84 | 8.76 | 2.4 | 2.92 | 1.88 | 0.191 | 0.334 |
| Temperate deciduous forest | 17.4 | 20.13 | 14.67 | 3.3 | 4.43 | 2.17 | 0.108 | 0.302 |
| Temperate evergreen forest | 14.5 | 15.98 | 13.02 | 3.6 | 4.39 | 2.81 | 0.176 | 0.337 |
| Temperate forest |  |  |  |  |  |  | 0.142 | 0.320 |
| Tropical deciduous forest | 15.8 | 19.15 | 12.45 | 7.4 | 9.16 | 5.64 | 0.295 | 0.736 |
| Tropical evergreen forest | 18.6 | 22.0 | 15.20 | 5.4 | 6.51 | 4.29 | 0.195 | 0.428 |
| Tropical forest |  |  |  |  |  |  | 0.245 | 0.582 |

† calculated from dataset of Table 3 in Jobbágy & Jackson (2000); $SC_{\text{density}} = (C_{\text{density}} \text{ in } 100 - 200 \text{ cm soil}) / (C_{\text{density}} \text{ in } 0 - 100 \text{ cm soil})$ .

**Table S6.** Global pattern of the proportions of forest plots in which soil porosity (SP) are lower than the estimated soil porosity in pre-developed forests (SP_0_).

| Biome | Plots with SP < = SP<sub>0</sub> | Total plots | Proportion (%) |
| --- | --- | --- | --- |
| Boreal | 11 | 41 | 26.8 |
| Temperate | 122 | 778 | 15.7 |
| Tropical | 31 | 142 | 21.8 |
| Total | 164 | 961 | 17.1 |
| Average |  |  | 21.4 |

**Table S7.**
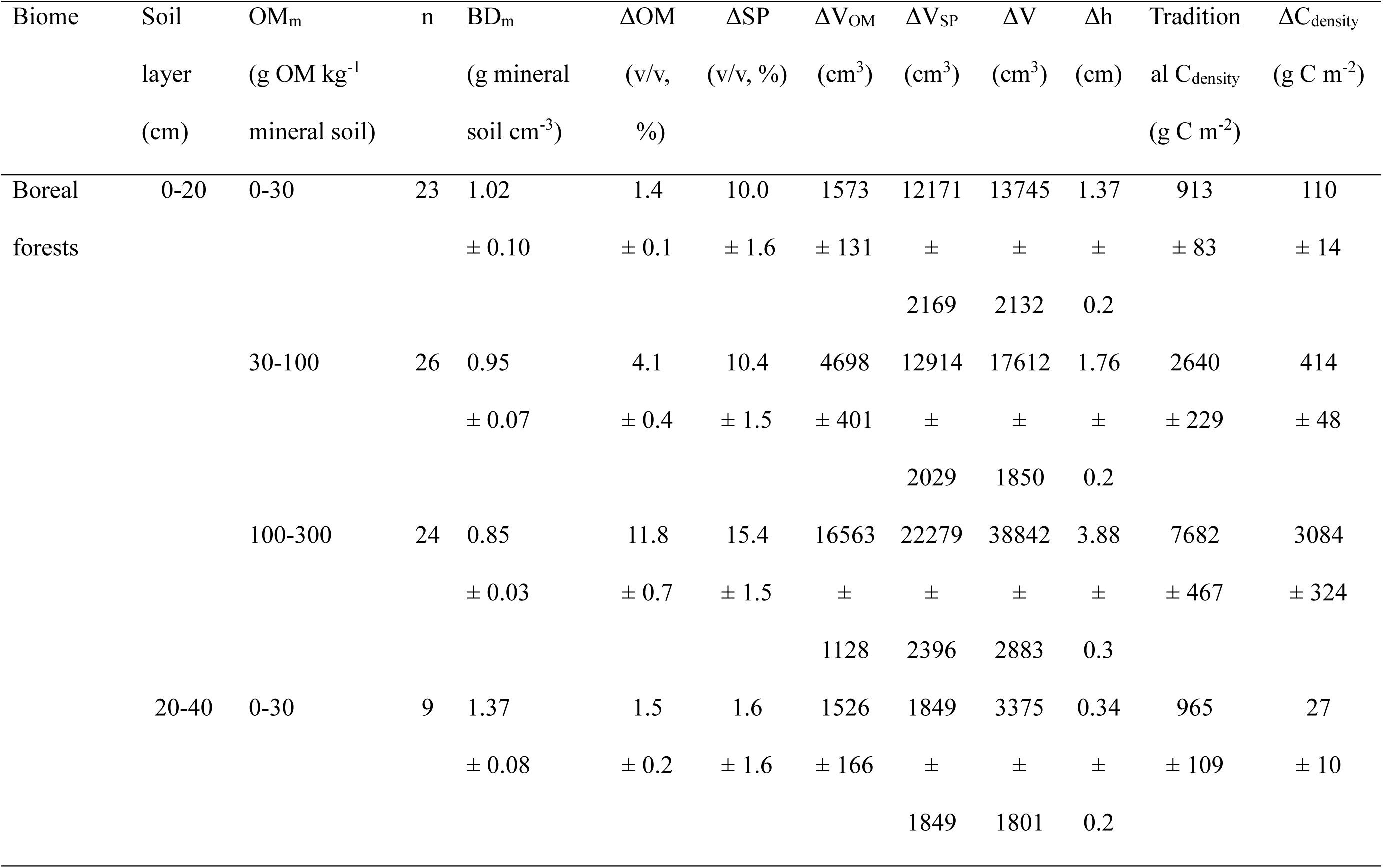
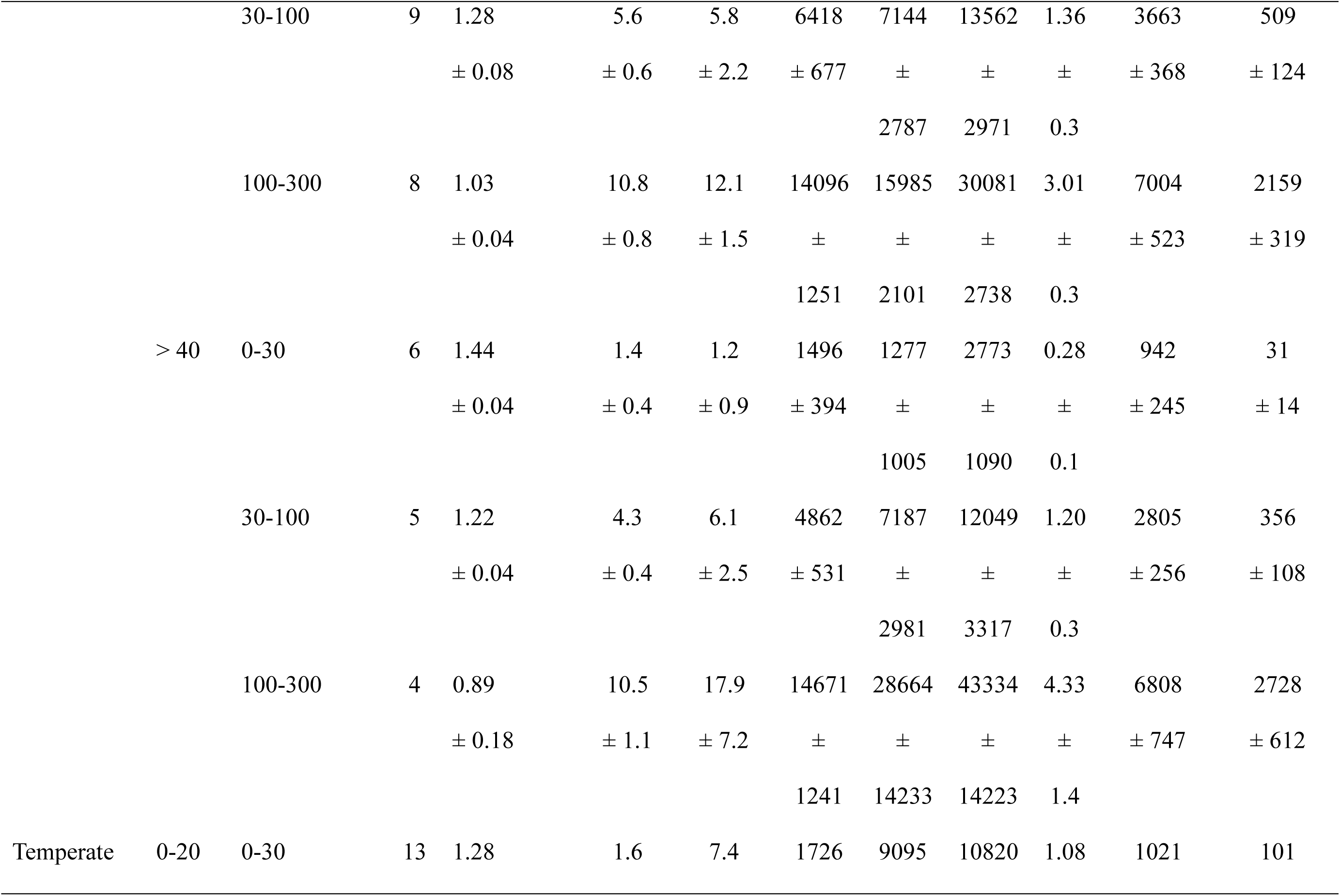
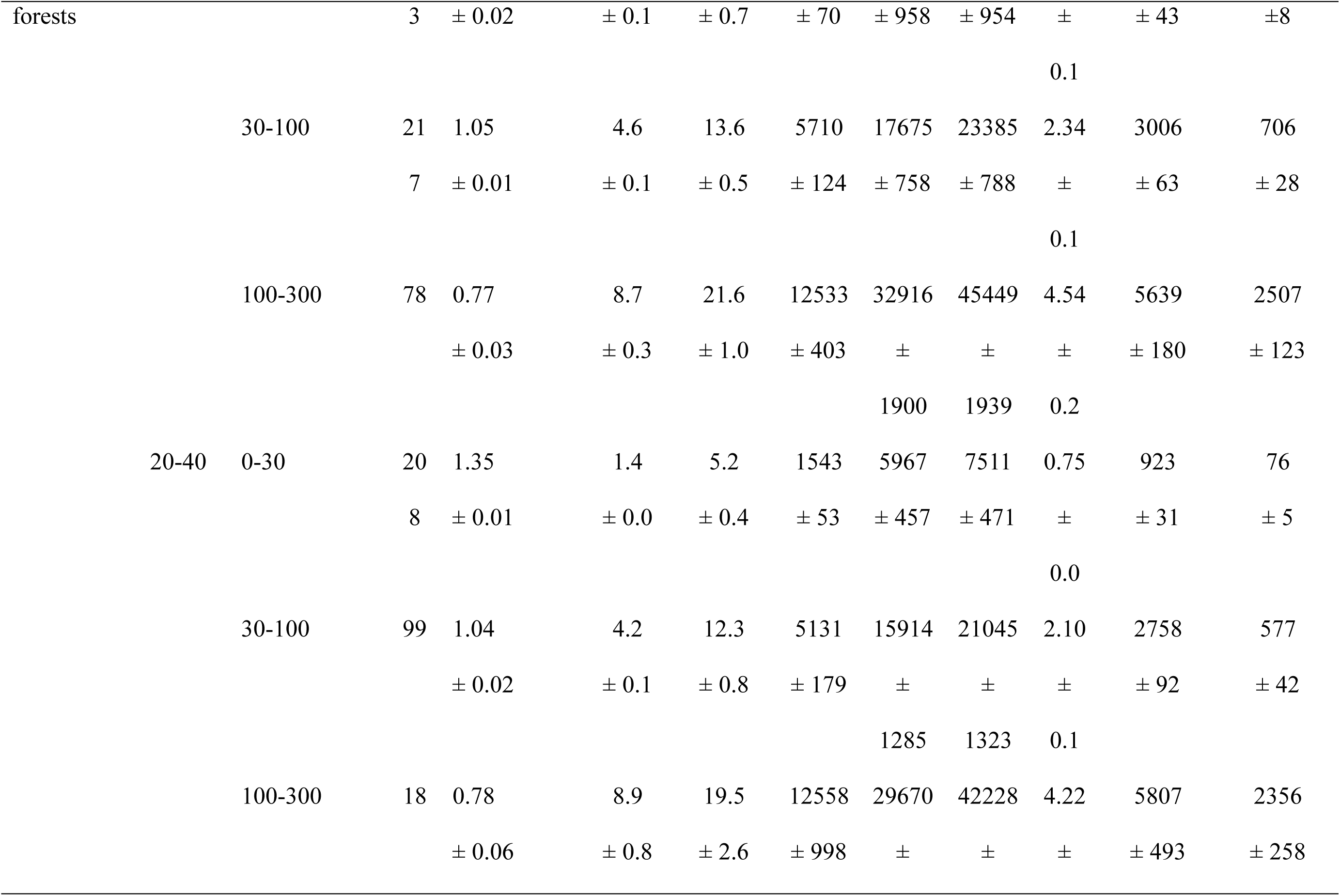
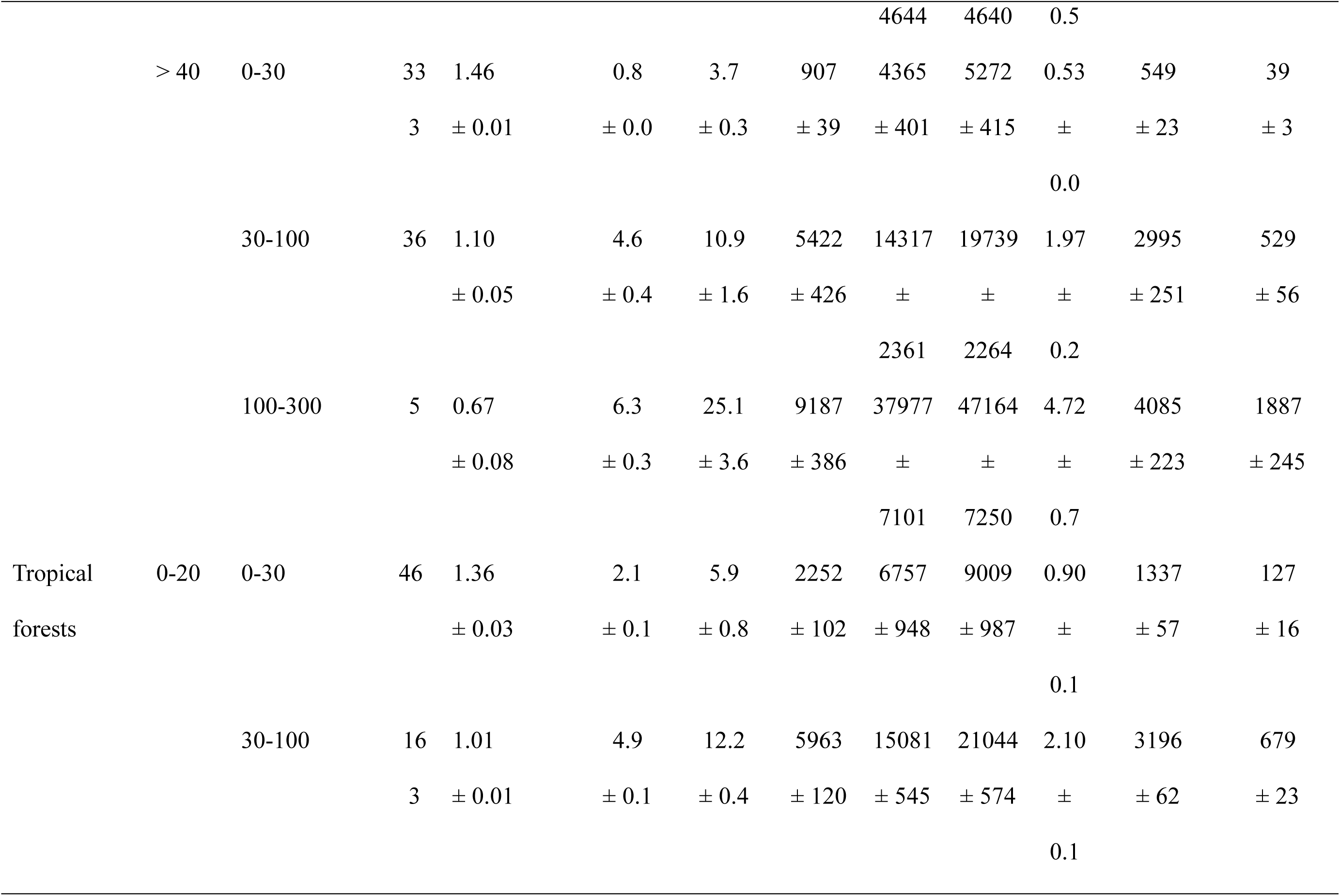
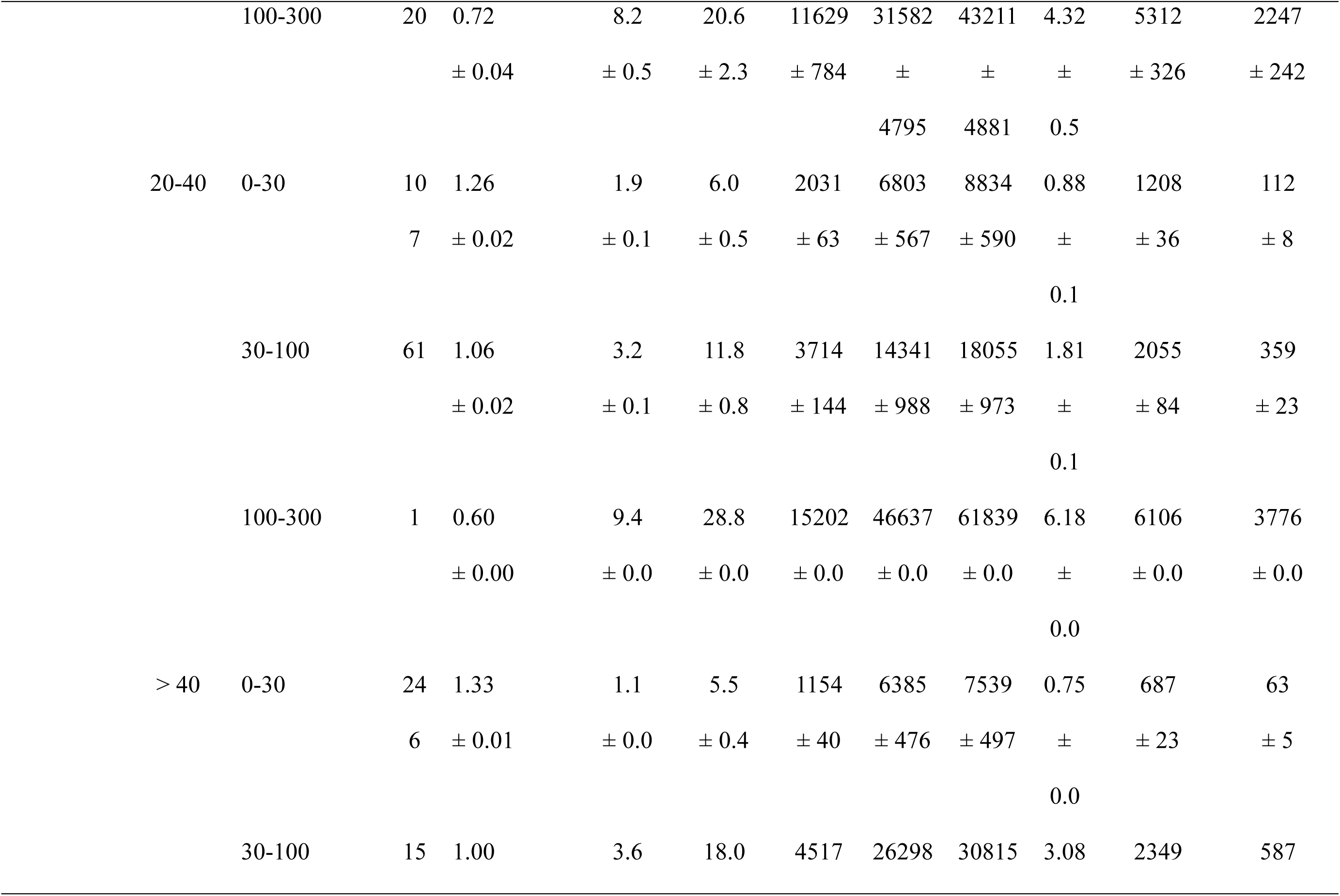
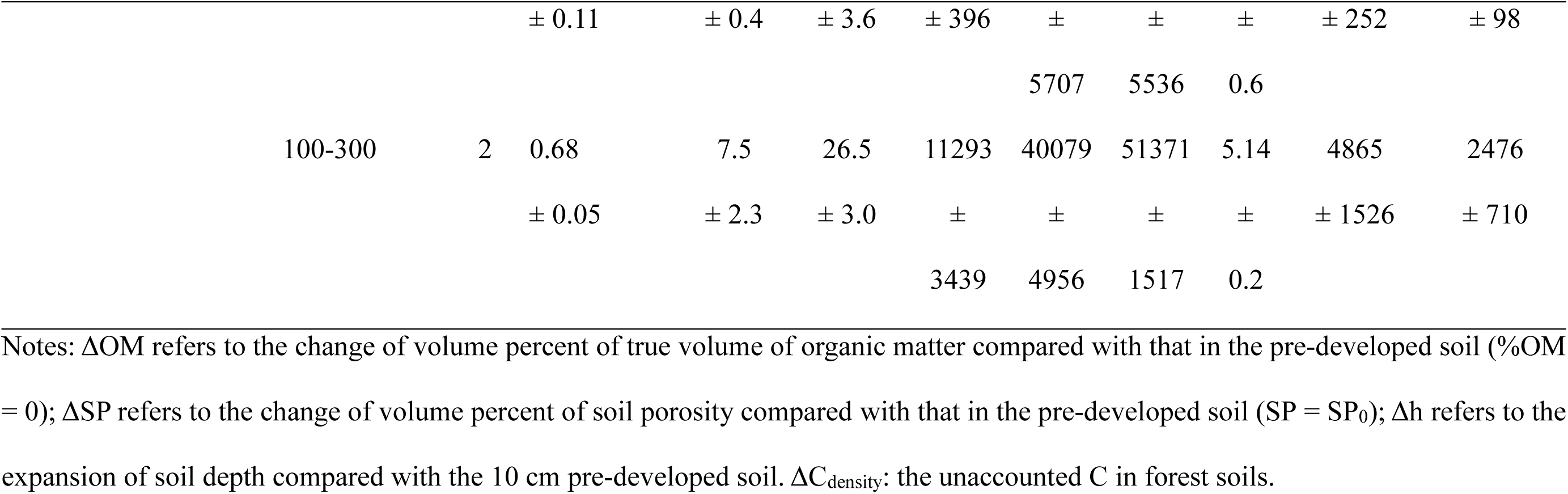
Theoretical change patterns of soil volume and unaccounted SOC in the standardized 10 cm mineral soil across biomes.

